# Changes in gene expression shift and switch genetic interactions

**DOI:** 10.1101/578419

**Authors:** Xianghua Li, Jasna Lalic, Pablo Baeza-Centurion, Riddhiman Dhar, Ben Lehner

## Abstract

An important goal in disease genetics and evolutionary biology is to understand how mutations combine to alter phenotypes and fitness. Non-additive interactions between mutations occur extensively and change across conditions, cell types, and species, making genetic prediction a difficult challenge. To understand the reasons for this, we reduced the problem to a minimal system where we combined mutations in a single protein performing a single function (a transcriptional repressor inhibiting a target gene). Even in this minimal system, a change in gene expression altered both the strength and type of genetic interactions. These seemingly complicated changes could, however, be predicted by a mathematical model that propagates the effects of mutations on protein folding to the cellular phenotype. We show that similar changes will be observed for many genes. These results provide fundamental insights into genotype-phenotype maps and illustrate how changes in genetic interactions can be predicted using hierarchical mechanistic models.

**One sentence Summary:** Deep mutagenesis of the lambda repressor reveals that changes in gene expression will alter the strength and direction of genetic interactions between mutations in many genes.

**Highlights:**

- Deep mutagenesis of the lambda repressor at two expression levels reveals extensive changes in mutational effects and genetic interactions
- Genetic interactions can switch from positive (suppressive) to negative (enhancing) as the expression of a gene changes
- A mathematical model that propagates the effects of mutations on protein folding to the cellular phenotype accurately predicts changes in mutational effects and interactions
- Changes in expression will alter mutational effects and interactions for many genes
- For some genes, perfect mechanistic models will never be able to predict how mutations of known effect combine without measurements of intermediate phenotypes

## Introduction

To interpret personal genomes, make accurate genetic predictions, and understand evolution we need to be able to predict the effects of mutations and also to understand how they combine (interact). Large-scale projects (Costanzo et al., 2016) and deep mutagenesis (Bank et al., 2015; Diss and Lehner, 2018; Domingo et al., 2018; Fowler et al., 2010; Hietpas et al., 2011; Melamed et al., 2013; Olson et al., 2014; Sarkisyan et al., 2016) have revealed that mutations frequently interact non-additively, which makes accurate genetic prediction a difficult challenge (Lehner, 2011).

Genetic (epistatic) interactions between gene deletions and loss-of-function alleles have been mapped genome-wide in budding yeast, revealing that both pairwise (Costanzo et al., 2016) and higher order (Dowell et al., 2010; Kuzmin et al., 2018) epistasis are widespread. Similarly, epistasis is widely detected when combining all possible pairs of mutations between two different proteins (Diss and Lehner, 2018), between natural genetic variants (Brem et al., 2005; Taylor et al., 2016) and between mutations selected during adaptation to new environments (Palmer et al., 2015; Weinreich et al., 2006). Systematic mutagenesis of individual proteins (Bank et al., 2015; Fowler et al., 2010; Hietpas et al., 2011; Melamed et al., 2013; Olson et al., 2014; Sarkisyan et al., 2016) and RNAs(Domingo et al., 2018; Li et al., 2016; Puchta et al., 2016) has also revealed widespread epistasis within individual macromolecules.

Moreover, comparisons across species (Dixon et al., 2008; Frost et al., 2012a; Roguev et al., 2008; Tischler et al., 2008), conditions (Bandyopadhyay et al., 2010; Díaz-Mejía et al., 2018a; Harrison et al., 2007) and cell types (Park and Lehner, 2015), have repeatedly found that genetic interactions are plastic, changing in different cells and conditions. This plasticity has important clinical implications for both evolution and genetic disease. For example, a ‘synthetic lethal’ genetic interaction between a cancer driver mutation and a drug or gene inhibition that can be exploited to specifically kill tumour cells of one type often proves ineffective in other cell types that carry the same driver mutation (Ashworth et al., 2011).

Why is this? Why do both the effects of mutations and genetic interactions change across conditions, cell types and species? Comparing between any two cell types, environmental conditions or species, there are typically thousands of molecular differences such as changes in gene expression, making this a difficult question to address. We reasoned that one way to address this question would be to ask it in a minimal system in which we could quantify the effects of mutations and how mutations interact and then test how these effects and interactions change in response to a simple perturbation of the cellular state. One of the simplest perturbations to make to a system is to change the expression level of a single gene, for example the expression level of the gene that is being mutated (Castel et al., 2018).

The phage lambda repressor (CI) is one of the best characterized proteins, serving as a paradigm for both gene regulation (Ptashne, 2004) and quantitative biology (Ackers et al., 1982; Igler et al., 2018; Lagator et al., 2017). The detailed and quantitative understanding of how this protein functions makes it an ideal system in which to address our question of how mutational effects and the interactions between mutations change when a system is perturbed.

Here we show that, even in this minimal system, the effects of individual mutations and the interactions between mutations change extensively when the expression level of the mutated gene is altered. Indeed we show that even a simple perturbation can result in the interactions between mutations changing in sign, flipping between positive (suppressive) and negative (enhancing) epistasis. We show that these seemingly complicated changes can be both understood and predicted using a mathematical model that propagates the effects of mutations on protein folding to the cellular phenotype. More generally, changes in gene expression will alter the effects of mutations and how they interact whenever the relationship between expression and a phenotype is nonlinear. Given that this is the case for most genes, shifts and switches in the interactions between mutations should be widely expected when the expression level of a gene changes.

## Results

### Deep mutagenesis of the lambda repressor at two expression levels

We used ‘doped’ oligonucleotide synthesis to introduce random mutations into the 59 amino acid helix-turn-helix DNA-binding domain of CI, and quantified the ability of each genotype to repress expression of a fluorescent protein (GFP) from the PR (Promoter R) target promoter by fluorescence-activated cell sorting into near neutral (Output1) and partially detrimental (Output2) bins and deep sequencing (Figure 1A – C). CI was expressed at a level similar to that observed in phage lysogens (Ptashne, 2004) (see STAR Methods). We quantified both the effects of single mutants and the genetic interactions between pairs of mutations. We then repeated the experiment expressing CI at a higher expression level and re-quantified the mutation effects and genetic interactions. The effects of wild type, 18 single and 4 double mutants when measured individually were highly correlated with their effects quantified in the pooled assay by deep sequencing at both expression levels (Figures 1D, rho=0.87, P<2e-16, n= 46; rho=0.82 and rho=0.71 respectively for low and high CI expression conditions, n=23, S1).

**Figure 1.**
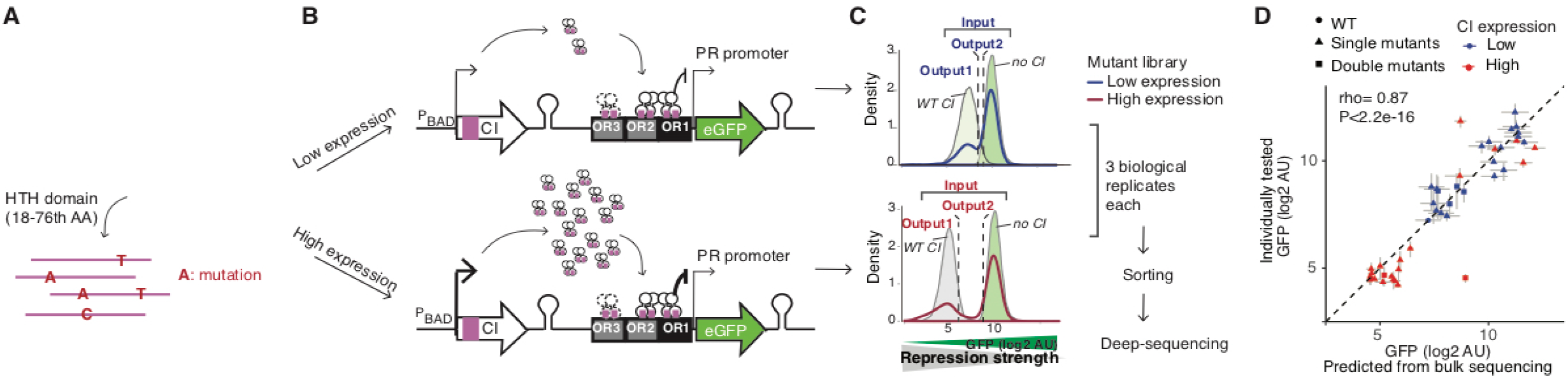
Deep mutagenesis of the lambda repressor CI DNA-binding domain at two expression levels (see also Figure S1). (A) CI-operator complex with mutated HTH domain in magenta. (B) Experimental design. (C) Distribution of GFP target gene expression when the mutant library is expressed at high and low levels and when WT or no CI is expressed. Sorted populations with GFP level similar to the wild-type population (Output1), and population with intermediate GFP level (Output2) were collected for deep sequencing. (D) Correlation of target gene expression estimated by deep sequencing with target gene expression individually quantified for wild type, 22 single and double mutants at low and high expression levels. Error bars denote standard error of the mean from four (y-axis) and three (x-axis) biological replicates.

At both expression levels, the single (Figure 2A, n=351) and double amino acid-change mutants (Figure 2B, n=468) had a bimodal distribution of target gene expression levels, with the low and high modes centred on the phenotypes observed for synonymous and premature stop codon-containing genotypes, respectively (Figure 2A, B). These bimodal distributions of mutational effects are consistent with observations for many different proteins (Diss and Lehner, 2018; Hietpas et al., 2011; Jiang et al., 2013; Melamed et al., 2013; Olson et al., 2014; Sarkisyan et al., 2016; Starr et al., 2018; Wylie and Shakhnovich, 2011), as is the shifted distribution of double mutant phenotypes towards higher expression of the target gene (i.e. reduced activity (Diss and Lehner, 2018; Sarkisyan et al., 2016)) (Figure 2A, B). Also consistent with previous deep mutagenesis datasets (Araya et al., 2012; Melamed et al., 2013; Olson et al., 2014; Sarkisyan et al., 2016), mutations in the core residues of the protein were more detrimental (reduced repression of the target gene) than mutations in solvent-exposed residues (Figures 2C, S2). Mutations in residues contacting DNA were also more detrimental than mutations in solvent-exposed residues (Figures 2C, S2). As expected, mutations to less similar amino acids were also more detrimental, as were mutations predicted to reduce the free energy of protein folding or DNA-binding (Figure S2C – F). Mutations to less hydrophobic amino acids were detrimental in the core and mutations that introduced a negative charge were detrimental at positions that contact DNA (Figure S2G – J).

**Figure 2.**
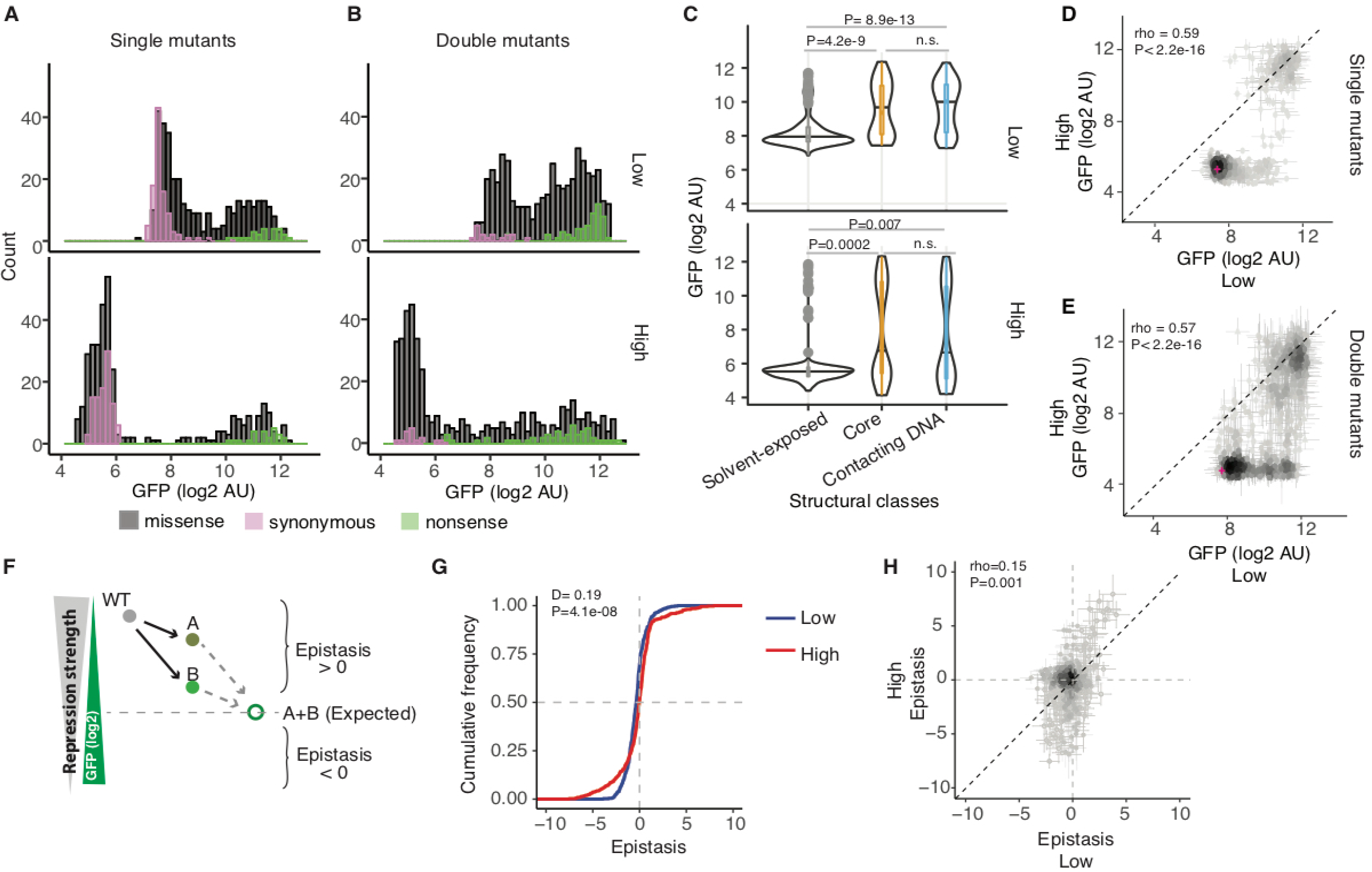
Comparison of mutational effects and genetic interactions at two expression levels (see also Figure S2). (A and B) Histogram of the mean mutational effects of single (n=351) (A) and double (n=468) (B) missense amino acid variants together with synonymous (n=114 for single, n=37 for double) and nonsense (n=21 for single, n=47 for double) variants. (C) Effects of single mutants in different structural regions. Classes compared using Wilcoxon Rank Sum test. (D and E) Comparison of mean mutational effects at the two expression levels. (F)Log-additive definition of epistasis. (G) Cumulative distributions of mean epistasis scores at the two expression levels (n=468). Distributions compared using two-sample Kolmogorov–Smirnov test. (H) Mean epistasis scores at the two expression levels. Error bars in (D, E, H) denote standard error of the mean.

### Mutation effects change non-linearly with a change in expression

Comparing the expression of the target gene when the same single (Figure 2D) or double (Figure 2E) mutant genotypes were expressed at high and low levels revealed a nonlinear relationship, with four main classes of genotypes: (1) genotypes with little effect at either high or low expression (~42% of single mutants), (2) genotypes having little effect at high expression but detrimental effects at low expression (~26% of single mutants), (3) genotypes that are partially detrimental at high expression but behave similarly to null alleles at low expression (~5% of single mutants), and (4) genotypes that behave similar to null alleles at both expression levels (~20% of single mutants). This ‘unmasking’ of detrimental mutation effects at low expression levels has been previously observed for mutations in a region of yeast Hsp90 (Jiang et al., 2013) and also for human disease-causing variants (Castel et al., 2018).

### Changing expression alters how mutations interact

We quantified epistasis between pairs of mutations as the difference between the observed and expected phenotypes based on a log additive model (Boucher et al., 2016). A positive epistatic interaction means that repression of the target gene by the double mutant is greater than expected and a negative interaction means that it is less than expected (Figure 2F). The distribution of epistasis scores differed between the two expression levels of the protein, with more strong positive and negative interactions at high expression (Figure 2G, two-sample Kolmogorov-Smirnov Test P=4.1e-8, D=0.19, n=468). Furthermore, epistasis scores of the same pairs of mutations at the two protein expression levels correlated only weakly (Figure 2H, rho=0.15, P=0.001,n=468). Plotting epistasis against the expected double mutational effects revealed systematic trends in the data (Figure S5A). Whereas double mutants with high expected target gene expression tended to interact positively at both low and high expression, double mutants with intermediate expected outcomes had stronger negative interactions at low expression, and double mutants with low expected target gene expression had stronger negative interactions at high expression (Figure S5A).

### A simple mathematical modelling predicts changes in mutational effects and interactions

What accounts for these systematic patterns of epistasis and also their dependence on expression level? To address this, we turned to a previously published quantitative model of repression of the PR promoter by CI (Ackers et al., 1982) (Figure S3A). Briefly, the model describes the probability of CI repressing the expression of the target gene as a function of the CI concentration (Figure 3A, C). We first mapped each single mutant’s effect from the target gene expression level to the concentration of active CI. We then extended this model to include the effects of mutations on the folding of CI and estimated changes in the free energy of folding for each single mutant (see STAR Methods). To predict the CI concentration and the resulting expression of the GFP target gene for each double mutant, we summed the change in free energy for each single mutant and then mapped the total free energy to a change in protein folding and concentration, which was in turn mapped to altered repression of the target gene (Figure S3B, C). We compared the behaviour of the full model (Figure 3B – E) to that of models that only considered protein folding (Figure S3D) or only repression of the target gene by CI (Figure S3E).

**Figure 3.**
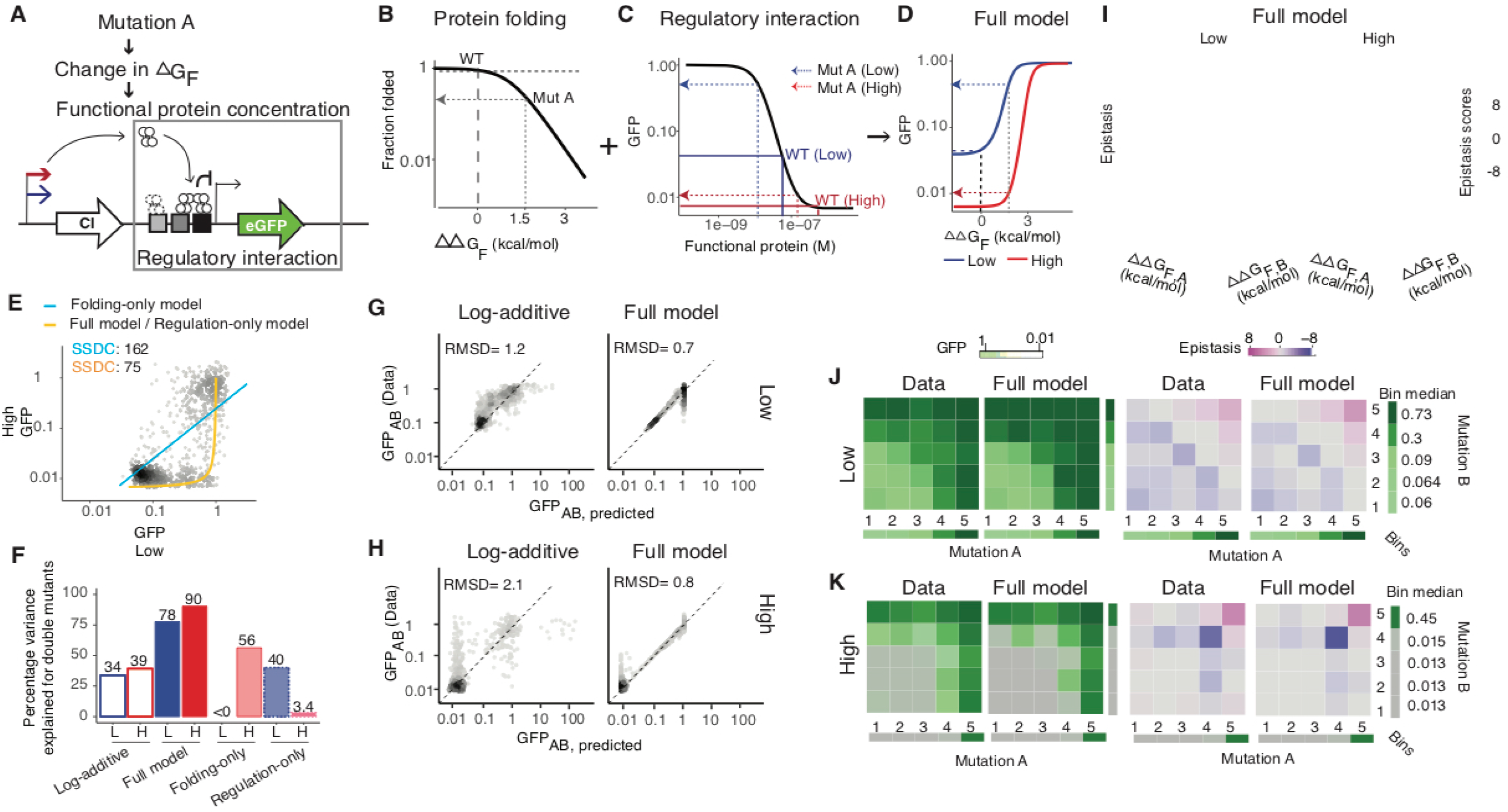
Combined model of protein folding and regulatory interaction predicts mutational effects and genetic interactions (see also Figures S3 – 5). (A) Mutations alter the free energy of protein folding (ΔG_F_) and so protein concentration and repression of the target gene. (B – D) Relationships between change in folding energy (ΔΔG_F_) and the fraction of folded protein (B), protein concentration and target gene expression (C), and change in folding energy (ΔΔG_F_) and target gene expression at low (blue) and high (red) expression (D). The effect of an example mutation (A) is indicated in each graph. (E) Regulatory interaction-only but not the protein folding-only model predicts the inverse relationship between target gene expression at the two expression levels. SSDC: sum of the squared distance from the curve. (F) Percentage of variance explained for double mutations for each model. ‘L’ indicates low expression and ‘H’ indicates high expression of CI. (G and H) Observed vs. predicted target gene expression for the log-additive and full folding + regulation model at low (G) and high (H) CI expression. RMSD: root-mean-square-deviation between the predicted and observed data. I, Predicted epistasis at low and high expression for the full model. (J and K) Model-predicted and experimentally-observed target gene expression and epistasis when combining mutants at low (J) and high (K) expression. Mutations were ordered by their effects into 5 equally-populated bins and the median target gene expression and epistasis plotted for each bin combination.

Both the full model and the transcription regulation-only model correctly predict the shape of the relationship between mutation effects at low and high expression (Figure 3E, n=819). However, only the full model provides good prediction of the phenotypes of double mutants from the phenotypes of the single mutants (Figure 3F). The full model (Figure 3G, H), but not the folding-only or regulation-only models (Figure S4), also captures the systematic trends in how mutations combine at both low and high expression.

### Nonlinear concentration-phenotype relationships cause expression-dependent epistasis

Inspection of the model reveals that it is the nonlinear relationship (Otwinowski et al., 2018) between protein concentration and target gene repression that causes the concentration-dependence of both mutational effects and genetic interactions (Figure 3D, I). Each mutation has a fixed effect on the free energy of protein folding (Figure 3B). When combining two mutations, the changes in free energy are summed and so alter the fraction of folded protein according to the nonlinear relationship in Figure 3B. However, because the relationship between protein concentration and target gene expression is also nonlinear, the same change in protein concentration can lead to a different change in target gene expression depending upon the starting protein concentration (Figure 3B – D). The nonlinear relationship between protein concentration and target gene expression therefore transforms the concentration-independent effects of mutations on protein folding (Figure 3B) into concentration-dependent changes in target gene expression (Figure 3C, D), resulting in concentration-dependent epistasis (Figures 3I – K, S5).

### Changes in gene expression reverse the sign of genetic interactions

Comparing how mutations combine at different expression levels in the full model revealed that changes in expression not only alter the magnitude of genetic interactions but can also switch their sign (between positive and negative interactions, Figure 4A, B). Re-analysis of the experimental data validated this prediction, with mutations in the regime predicted by the model switching from positive to negative epistasis as the expression level increased (Figures 4C, S6). In other words, genetic interactions that are suppressive at one expression level can become enhancing at another expression level (Figure 4D,E). Our model and data therefore show for the first time that changes in expression can alter both the strength and the type of epistasis between mutations.

**Figure 4.**
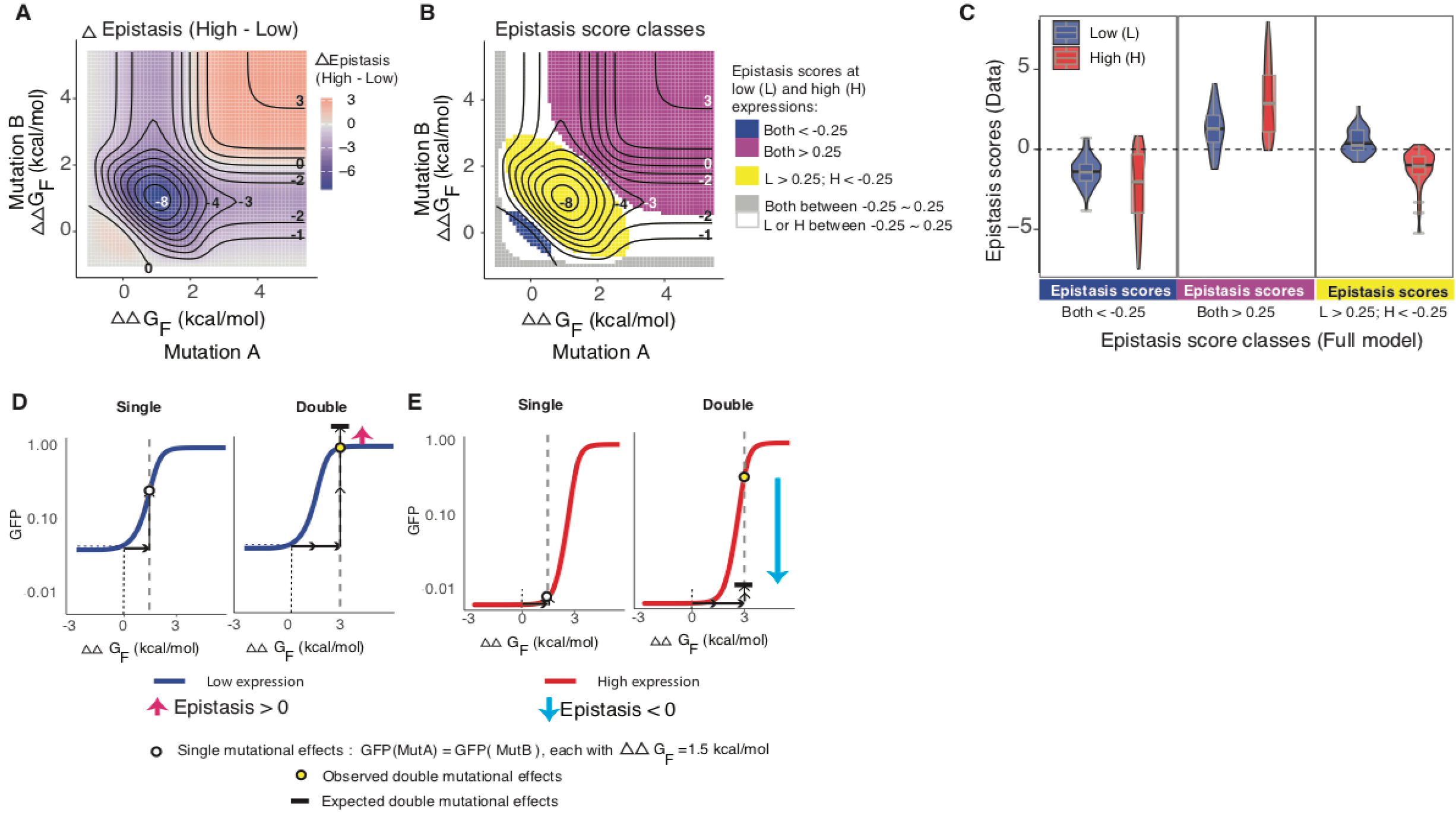
Changes in expression alter both the strength and sign of epistasis (See also Figure S6). (A and B) Changes in epistasis strength (A) and class (B) between low and high expression predicted by the model. (C) Experimentally-determined epistasis scores for double mutants with the indicated model-predicted epistasis scores. (D and E) The same pair of mutations can interact positively at low expression (D) and negatively at high expression (E).

### Changes in gene expression will alter genetic interactions for many genes

To what extent should we expect these conclusions to apply to other genes? Mutational effects and genetic interactions will be expression-level dependent whenever the relationship between expression and a phenotype is nonlinear. Such nonlinear expression-fitness functions are indeed very common in biology because of the abundance of cooperation, competition, and feedback, with nonlinear functions used to model almost all aspects of cell biology (Bhaskaran et al., 2015). Moreover, the relationship between expression level and fitness (growth rate) has been systematically quantified for 81 yeast genes and, for all genes sensitive to a change in expression in the tested conditions, the expression-fitness function was nonlinear (Keren et al., 2016).

We quantified epistasis and its sensitivity to concentration changes in three of the most common expression-fitness functions of yeast genes (Keren et al., 2016). For many yeast genes, fitness increases as a concave function as their expression is increased from zero to a fitness plateau close to the wild-type expression level (Figure 5A). For these genes, epistasis changes in magnitude but not sign as the expression level changes (Figures 5C, E, G, S7). Similar results are seen for genes where fitness decreases as a concave function as expression is increased (Figure 5). Multiple genes in yeast have a ‘peaked’ expression-fitness landscape(Keren et al., 2016). For these genes epistasis can change substantially and also switch in sign as the expression level changes because of the non-monotonic relationship between the free energy of protein folding and fitness (Figure 5B, D, F, H).

**Figure 5.**
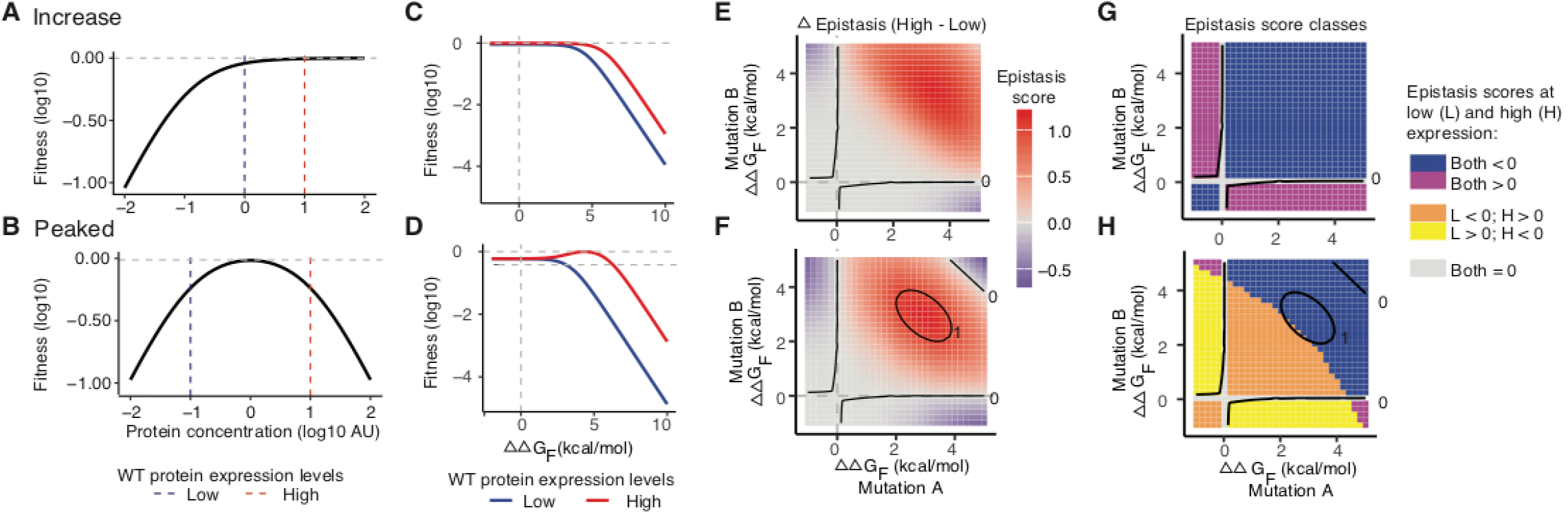
Other common expression-fitness functions generate concentration-dependent genetic interactions (See also Figure S7). (A and B) Two common expression-fitness functions in budding yeast. (C and D) Relationship between change in free energy of protein folding and fitness for these functions, (E and F) Change in epistasis magnitude between high and low expression. (G and H) Change in epistasis sign between high and low expression.

### Non-monotonic expression-phenotype relationships result in ambiguous genetic prediction

Finally, analysing how mutations combine in genes with different expression-fitness functions we realised that for some genes accurate predictions for how mutations combine will never be possible, even with a perfect mechanistic model. Specifically, when there is a non-monotonic relationship between the expression level and a phenotype, the same observed phenotype for a single mutant can map to two or more different free energies of protein folding, leading to multiple possible double mutant phenotype predictions for each mutation pair (Figures 6, S8). For these genes, even a perfect mechanistic model is therefore insufficient to predict how mutations of precisely measured effects combine to alter a phenotype. In such cases it will always be necessary to make additional measurements – for example of intermediate phenotypes such as protein concentrations – to predict how two mutations will combine to alter a phenotype.

**Figure 6.**
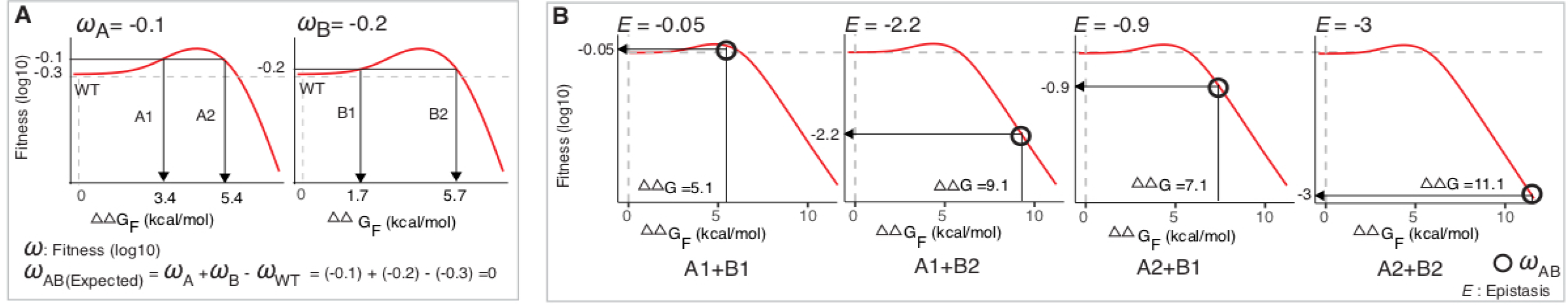
Unpredictable double mutant phenotypes (See also Figure S8). (A) For ‘peaked’ expression-fitness functions such as that shown in Figure 5B, the same change in fitness can be caused by two different changes in folding free energy. (B) For a pair of single mutant phenotypes there can therefore be up to 4 possible double mutant outcomes.

## Discussion

Non-additive interactions between mutations greatly complicate genotype-phenotype maps and so make genetic prediction a difficult challenge. That the interactions between mutations change across conditions, for example between different types of cancer (Park and Lehner, 2015), further complicates this. To better understand the plasticity of both mutational effects and genetic interactions, we studied the question in a very well-understood minimal system where we could quantify how both the effects of mutations and their interactions change in response to a simple perturbation. We chose to use the lambda repressor as a model system because it is one of the best-understood regulatory proteins in biology and because there are very well established and accurate mathematical models that describe its regulatory activity (Ackers et al., 1982).

For the lambda repressor, we found that a change in expression altered both the effects of individual mutations and how these mutations combined. Moreover, we found that changes in expression altered both the strength and the type of interactions between mutations, with many mutations switching from positive (suppressive) to negative (enhancing) epistasis at different expression levels.

Although our experimental work focussed on the lambda repressor, by analysing other common expression-fitness functions, we have shown that our conclusions will widely apply to many genes. Indeed changes in expression will transform the effects of mutations and their interactions whenever the relationship between expression and a phenotype is nonlinear. In yeast, where expression-fitness functions have been systematically quantified (Keren et al., 2016), this is normally the case: for most genes the growth rate of the organism does not depend in a linear way on the gene’s expression level. For many genes, therefore, changes in expression alone will drive changes in mutational effects and genetic interactions. Thus we should expect that genetic interactions will change extensively across conditions and cell types in an animal, as well as between individuals in a population and between different species. Analyses of genetic interactions across conditions (Bandyopadhyay et al., 2010; Díaz-Mejía et al., 2018b; Harrison et al., 2007; Onge et al., 2007), cell types (Ashworth et al., 2011; Park and Lehner, 2015), and species (Dixon et al., 2008; Frost et al., 2012b; Roguev et al., 2008; Tischler et al., 2008) are highly consistent with this.

Changes in genetic interactions are highly relevant to both agriculture (Soyk et al., 2017) and human genetic disease. For example, dynamic epistasis may contribute to the tissue-specificity of human disease mutations as well as the cancer type-specificity of interactions between cancer driver mutations (Park and Lehner, 2015). Moreover, the success of synthetic lethal strategies to specifically kill target cells depends on the stability of these interactions. Many examples now exist of synthetic lethal gene perturbations that are effective in one cancer cell type but ineffective in other cell types, and the most successful targets will be interactions that are very stable across individuals and perturbations (Ashworth et al., 2011; Park and Lehner, 2015).

Finally, the plasticity of epistasis will also need to be incorporated into evolutionary models. Epistasis is a strong determinant of evolutionary paths (Breen et al., 2012; Starr and Thornton, 2016). The plasticity of epistasis caused by changes in the expression level suggests that the accessible and most likely evolutionary paths will change over time as the expression level of a gene is altered.

Very importantly, we found that the seemingly complicated shifts and switches in genetic interactions as the expression level of the lambda repressor changed could be both understood and accurately predicted using a hierarchical mechanistic model that propagates the effects of mutations on the free energy of protein folding to the cellular phenotype. Considering just the effects of mutations on protein folding or just how the repressor regulates gene expression could not account for the changes in interactions. We envisage that such multi-step models that propagate the effects of mutations on protein stability to higher-level phenotypes may prove generally useful for genetic prediction and for understanding how mutations combine to alter phenotypes, including in human disease.

In our analyses we only considered the effects of mutations that alter the free energy of protein folding. Altered protein stability is likely to be by far the most common affect of amino acid changing mutations (Tokuriki and Tawfik, 2009a). However, subsets of mutations will have additional effects, for example altering the affinity and kinetics of molecular interactions. In future work it will be important to study how mutations with different molecular effects interact with each other, as well as with mutations that affect stability and with changes in expression. Our model also makes the assumption that the effects of mutations on protein stability are independent of the expression level but this may sometimes not be the case, for example because of chaperone titration (Tokuriki and Tawfik, 2009b) or interactions with other molecules (Bridgham et al., 2009; Diss and Lehner, 2018). Concentration-dependent changes in the effects of mutations on protein stability will lead to further shifts in mutational effects and genetic interactions as a gene’s expression changes.

Finally, although we found that a hierarchical model provided accurate genetic prediction for the lambda repressor, we also realised that there are cases where such a mechanistic model will fail to accurately predict how mutations combine to alter phenotypes. Specifically, when there is a non-monotonic relationship between the concentration of a protein and a phenotype, it is sometimes not possible to predict how two mutations will combine, even with a detailed mechanistic model. This is because some phenotypes map to two or more possible changes in protein concentration and so to multiple changes in the free energy of protein folding. Without additional measurements it is not possible to tell which of the underlying changes is causing the phenotype. This results in multiple possible outcomes when mutations of known phenotypic effect are combined. In these cases, additional measurements of intermediate phenotypes such as protein concentrations will always be required for accurate genetic prediction.

## Acknowledgements

We thank members of the Lehner lab and J. Ren for comments on the manuscript. This work was supported by a European Research Council (ERC) Consolidator grant (616434), the Spanish Ministry of Economy and Competitiveness (BFU2017-89488-P and SEV-2012-0208), the Bettencourt Schueller Foundation, Agencia de Gestio d’Ajuts Universitaris i de Recerca (AGAUR, 2017 SGR 1322.), and the CERCA Program/Generalitat de Catalunya. X. Li was supported in part by a fellowship from the Ramón Areces Foundation.

## Authors Contributions

X.L. performed all experiments, analyses, and modelling. J.L. built the plasmid construct pCIPR. X.L., J.L., R.D., and B.L. conceived the project. X.L. and B.L. designed the project and interpreted the data, X.L., P.B-C., and B.L. wrote the manuscript.

---

**Figure S1. Reproducibility of mutational effects between biological replicates. Related to Figure 1.**

(A and B) Spearman correlations of mutational effects among three biological replicates for low (A) and high (B) CI expression.

(C) Comparisons of mutational effects between low and high expression level for 22 individually retested single and double mutants together with wild type. Error bars denote standard error of the mean.

(D) Density plots of GFP expression for the 22 individually re-tested single and double mutants at the two expression levels of CI.

**Figure S2. Mutational effects depend on both the chemical features of amino acid substitutions and the tertiary structural positions. Related to Figure 2.**

(A) Structure of CI dimer bound to an operator (PDB 3bdn). One monomer is shown as a ribbon and the other one with all its atoms shown as spheres. Only the mutagenized HTH domain is shown. Left panel is the structural classification of the residues. Middle and right panels show the positional median z-scores of GFP expression levels after subtracting wild type z-scores at the two expression levels of CI. Z-scores rather than absolute GFP expression levels are shown here to compare positional sensitivity to mutations at two expressions of CI.

(B) Heatmaps of mean GFP expression for single mutations at the two expression levels. Amino acids are ordered based on their similarities, from top to bottom: hydrophobic aromatic (F,W,Y), hydrophobic nonpolar aliphatic (P,M,I,L,V,A,G), hydrophilic polar uncharged (C,S,T,N,Q), hydrophilic negatively charged (D,E) and hydrophilic positively charged (H,K,R). Wild type amino acids are shown as letters inside the heatmap.

(C and D) Target GFP expression compared to the amino acid substitution matrix scores (BLOSUM62) at low (C) and high (D) expression of CI.

(E and F) Target GFP expression compared to the FoldX-predicted changes in the folding energy of the protein (E) and protein-DNA binding (F) at the two expression levels. Linear regression lines for each structural class are shown with the shaded areas showing the permutation-based 95% confidence intervals for the fit. ns – not significant.

(G and H) Target gene expression compared to the change in the hydrophobicity at low (G) and high

(H) expression of CI.

(I and J) Target gene expression compared to to changes in the side chain charges at low (I) and high

(J) expression of CI.

**Figure S3.** Mathematical models. Related to Figure 3.

(A) Eight configuration states (CS) of the PR promoter.

(B) Obtaining functional protein concentration (panels b1,b4), fraction of folded protein (panels b2,b3), and change in folding energy (panels b2,b3) from GFP expression levels of a mutation at low expression of the protein.

(C) Scheme for predicting double mutants’ GFP expression levels from single mutants’ GFP expression levels based on different models.

(D) Folding-only model.

(E) Regulation-only model.

**Figure S4. Predictions of double mutants based on folding-only or regulation-only model. Related to Figure 3.**

(A and B) Observed versus predicted GFP expression levels for the folding-only (A) and regulation-only (B) models. RMSD: root-mean-square-deviation from the predicted to the observed data.

(C – F) Binned median target gene expression levels (C, E) and epistasis scores (D, F) for the folding-only (C, D) and regulation-only (E, F) models. Mutations were sorted into 5 equally populated bins by their single mutant phenotypes as in Figure 3J,K.

**Figure S5. Epistasis pattern predicted from different models. Related to Figure 3.**

(A – D) Epistasis versus GFP expression levels for observed data (A), predicted from full model (B), folding-only model (C) and regulation-only model (D).

(E – G) Epistasis scores at the two expression levels of CI protein for full model (E), folding-only model (F) and regulation-only model (G). Two-sample Kolmogorov–Smirnov test was performed for cumulative distributions of epistasis scores at the two expression levels of CI protein.

**Figure S6. Observed versus predicted expression level-dependent changes in epistasis. Related to Figure 4.**

(A) Histogram of the model-predicted epistasis score distributions at the two expression levels of the protein. The grey dotted lines mark the center bin with the epistasis score thresholds of −0.25 and 0.25; and the black dotted lines mark the center three bins with the epistasis score thresholds of −0.75 and 0.75.

(B and C) Distribution of the observed epistasis scores grouped by the model-predicted classes of epistasis scores, with classification threshold of −0.25 and 0.25.

(D and E) Distribution of the observed epistasis scores grouped by the model-predicted classes of epistasis scores, with two additional classification thresholds, between −0.75 and 0.75 (D) and between −0.1 and 0.1 (E). “L” - low expression and “H” - high expression.

**Figure S7.** Concentration-dependent genetic interactions in the yeast fitness landscape. Related to Figure 5.

(A – C) Concentration-dependent mutation effects and epistasis in a “decreasing” expression-fitness function(Keren et al., 2016).

(D – L) Concentration-dependent epistasis for three common expression-fitness functions with stable, marginally stable and unstable proteins.

**Figure S8.** Unpredictable double mutant phenotypes. Related to Figure 6.

(A) A measured fitness effect can be caused by two different changes in protein concentration in a ‘peaked’ fitness landscape when the WT protein is expressed at the fitness optimum.

(B) Only very small changes in fitness can be mapped to either increased or decreased fraction of folded protein, due to the limit of fraction of folded protein (maximum equals to 1). For example, a mutant with the fitness effect of −0.02 (ω_A_) can be caused by two different mutations (A1 and A2) that cause changes in the free energy of protein folding (ΔΔG_F,A1_ or ΔΔG_F,A2_) and so two different changes in protein concentration. In contrast, larger fitness changes can only be caused by one change in free energy of folding. For example, a mutant with a fitness effect of −0.05 (ω_B_) can be caused by either a 5-fold increase or decrease in the functional protein concentration. However, a 5-fold increase in concentration cannot be achieved by a change in folding because it would require more than 100% of the protein to be folded. Therefore, a mutant with a fitness effect of −0.05 can only be caused by a decrease in protein stability (mutant B1).

(C) Combining two mutations of known fitness can lead to two possible double mutant outcomes and either positive or negative epistasis. For the case of A2 + B1, mutant A2 is detrimental in the wild type background (ω_A2_=-0.02), but beneficial at the mutant B1 background (ω_A2B1_-ω_B1_= −0.02 –(−0.05)= 0.03). The interaction between mutant A2 and B1 is. Therefore an example of sign epistasis. The possible outcomes are up to 4 if the fitness landscape is not symmetrical.

**Figure S9.** Fluorescence-activated cell sorting (FACS). Related to STAR Methods.

(A and B) An example (High expression, replicate 3) of the gating strategy for FACS.C. FACS recordings from each biological replicate performed on different days. GFP_index is used to quantify variation in fluorescence readings between batches.

**Figure S10.** Protein quantification. Related to STAR Methods.

(A) Distribution of fluorescence signal of cells expressing C-terminal GFP-tagged CI at high and low expression levels.

(B) Fluorescence linearly correlates with the number of molecules of equivalent soluble fluorochrome (MESF) from GFP beads.

(C) Relative fold-change of soluble CI protein concentrations at high versus low expression levels. Error bars denote standard error of the mean.

**Figure S11.** Filtering of sequencing data. Related to STAR Methods.

(A and B) Sequencing data was filtered to only retain genotypes with at least 100 read counts (red line) in all three biological replicates for both low (A) and high (B) expression datasets. Each smooth scatter panel shows the relationship between enrichment scores (S_v,o1_ for Output1 and S_v,o2_ for Output2) and input read counts for each replicate. The top density plot shows the input count distribution for each replicate.

(C and D) Only variants with propagated mean enrichment score standard errors smaller than 1 (red line) were retained.

**Figure S12.** Converting enrichment scores to GFP expression. Related to STAR Methods.

(A – C) Relationship between GFP signals either with Output1 enrichment scores (A), with Output2 enrichment scores (B), or with transformed Output2 enrichment scores (C) for the individually tested variants (n=23).

(D and E) Relationship between Output1 and Output2 enrichment scores (D) or transformed Output2 enrichment scores (E) for all single nucleotide variants (n=531).

(F and G) Comparisons of individually tested mean GFP signals with the predicted mean GFP signals from Output1 and Output2 enrichment scores (F) or with Output1 and transformed Output2 enrichment scores (G) (n=23). All error bars denote standard error of the mean. RMSD: root-mean-square-deviation between the predicted and observed data, after averaging the replicates.

**Figure S13.** Correcting for technical biases. Related to STAR Methods.

(A) Relationship between predicted GFP expression for biological replicates for all single nucleotide variants (n=531) before (gray) and after (blue or red) transforming the replicate 1 and 3 data to the reference replicate 2 (see Methods).

(B and C) Density plot of GFP expression before (B) and after (C) correcting for technical biases by transforming replicates 1 and 3 to the reference replicate 2 for all single nucleotide variants (n=531).

(D) Smooth scatter showing the relationship between the mean GFP signal of all amino acid genotypes (n=888) before and after scaling to the detection range (see Supplementary Methods).

**Figure S14.** Mathematical modelling. Related to STAR Methods.

(A) Relationship between free CI dimer concentration and total CI concentration in the cell in Ackers’ model.

(B – D) Parameter search for the line intercept that best describes the relationship of GFP at low and high expression for the folding-only model. Dashed lines in (B) and (D) mark equal GFP level at the two expression levels. Solid lines in (B) mark the range of the intercepts searched for the best fit. Red dashed line in (C) shows the best fit (the smallest SSDC).

(E) Projection of individual data points from observed GFP expression levels at low and high CI expression to the model-predicted curve.

## STAR Methods

### CONTACT FOR REAGENT AND RESOURCE SHARING

Further information and requests for resources and reagents should be directed to and will be fulfilled by the Lead Contact, Ben Lehner

### EXPERIMENTAL MODEL AND SUBJECT DETAILS

#### Microbe strain and growth conditions

*E.coli* BW27783 MK01 strain (kindly provided by the M.Isalan lab), modified to homogenously express arabinose-induced genes (Kogenaru and Tans, 2014) was used to express the mutant library. A single colony of the *E.coli* BW27783 MK01 strain was picked Luria-Bertani (LB) agar plate, grown overnight at LB liquid medium supplemented with chloramphenicol to 14μg/ml concentration at 37°C. The 500μl overnight growth media with cells mixed with 500μl of 50% glycerol were stored at −80°C freezer. For experiments, cells were always grown at 37°C in LB liquid medium supplemented with appropriate antibiotics. For specific experimental growth conditions, please refer to the method details in the following section.

## METHOD DETAILS

### Experimental Methods

#### Mutant oligonucleotide library synthesis and amplification

A 250-nucleotide-long oligonucleotide library was synthesized by TriLink BioTechnologies. Library oligonucleotides contain a 177-nucleotide-long sequence of the CI Helix-Turn-Helix domain (52th-210th nucleotide bases, based on CI ORF GeneID:3827059), “doped” at each position with 0.4% of each of the three non-reference nucleotides. The “doped” region is flanked by invariant sequences corresponding to the wild type sequences of immediate upstream (36 nucleotide bases) and downstream (37 nucleotide bases) of the doped region and used as constant overhang regions for the PCR primers to bind. The designed oligonucleotide sequence is:

5’*CCATTAACACAAGAGCAGCTTGAGGACGCACGTCGCcttaaagcaatttatgaaaaaaagaaaaatgaacttggcttatcccaggaatctgtcgcagacaagatggggatggggcagtcaggcgttggtgctttatttaatggcatcaatgcattaaatgcttataacgccgcattgcttgcaaaaattctcaaagttagcgttgaagaatttAGCCCTTCAATCGCCAGAGAAATCTACGAGATGTATG* 3’

Upper case indicates the constant regions and lower case the “doped” sequence.

The ‘doped’ library was dissolved in 500ul MilliQ water as a stock solution, and 10μl of the stock solution was further diluted in 500ul of MilliQ water as a working solution. The working solution oligonucleotide concentration was estimated to be 390ng/μl based on NanoDrop (Thermofisher Scientific) measurement of ssDNA concentration. Next, the working solution ‘doped’ library was further diluted by a factor of 100, and a total of about 40ng was used as the template to synthesize the complementary strand as well as to be amplified. Polymerase chain reaction (PCR) was performed using Phusion high fidelity PCR kit (Thermo Scientific) with primers that bind to the constant regions of the ‘doped’ library oligonucleotide (Table S1). Each 50μl PCR reaction consisted of 10μl of the ‘doped’ library oligonucleotide as the template, 10μl of 5X Phusion HF reaction buffer, 1μl of 10mM dNTP (NEB), 2.5μl of 10μM forward and reverse primers each, 0.5μl of Phusion polymerase and 12.5μl MilliQ. PCR reactions followed the manufacturer’s instruction for a standard protocol.18 PCR cycles were performed to minimize incorporation of PCR errors to the library. PCRs were performed with annealing temperature at 55°C and extension at 72°C for 30 seconds. The fragment with the correct size (230 nucleotide bases) was visualized and retrieved using the 2% size-select E-gel purification system (Invitrogen). To achieve optimal ligation efficiency, the size-selected PCR fragment was further purified with the MiniElute PCR purification kit (QIAGEN) to remove excess salt. The Gibson assembly (GA) system was used to ligate the PCR fragments to the modified plasmid backbone (see below) following the standard GA protocol.

### Plasmid constructs and the expression of the mutant plasmid libraries

The CI open reading frame (GeneID:3827059) was cloned into the bacterial expression vector pBADM-11 (obtained from CRG biomolecular screening & protein technologies unit), between the arabinose-inducible promoter pBAD and three stop codons in all three reading frames (“tagttaagtga”), followed by the strong synthetic bidirectional terminator L3S2P21(Chen et al., 2013). The PR promoter (that overlaps with OR3, OR2 and OR1 repressor binding sites) followed by the RBS-GFP (LVA) ORF(Andersen et al., 1998) was cloned downstream of the L3S2P21 terminator. Three stop codons in all three reading frames (“tagttaagtga”) were cloned immediately downstream of the GFP ORF and upstream of the pBADM-11 intrinsic rrnB_T2 terminator.

The two plasmid constructs (pCIPR plasmids) used in our experiments—the construct expressing CI to a high concentration (pCIPR-High) and the construct expressing CI to a low concentration (pCIPR-Low)—differed in the DNA sequences between the predicted strongest ribosome binding sequence (RBS) and the ATG start codon of the CI gene. In pCIPR-High, the start codon is immediately after the RBS. In pCIPR-Low the start codon is 82 nucleotides downstream of the RBS.

The pCIPR-High and pCIPR-Low plasmids were linearized by removing the coding region of the CI helix-turn-helix motif (HTH) domain that contains the ‘doped’ sequence. The doped oligonucleotide library and the linearized plasmids were assembled using the GA system (master mix provided by CRG biomolecular screening & protein technologies unit) following the standard protocol. The assembly reactions were dialysed using 0.025um VSWP membrane filters (Merk Millipore Ltd) and electroporated into the high efficiency commercial NEB10β competent cells (NEB, C3020K). After recovery in 500μl Super Optimal broth with Catabolite repression (SOC) culture media at 37°C for one hour, an aliquot of the cells was plated on Luria-Bertani (LB) agar plate with 100μg/ml ampicillin to examine the transformation efficiency, and the rest was diluted 1 in 200 in fresh Luria-Bertani (LB) broth with 100μg/ml ampicillin for overnight growth. About 780,000 independent transformant colonies were obtained for the mutant plasmid library construction. Plasmids were purified using the Qiagen Midiprep kit (cat.12143) and the purified plasmids were then used as the mutant plasmid library.

#### Making highly efficient electro-competent cells

We chose the *E.coli* BW27783 MK01 strain (kindly provided by the Isalan lab)(Kogenaru and Tans, 2014), modified to homogenously express arabinose-induced genes, to express the mutant library. A single chloramphenicol-resistant colony of was picked into 4ml LB medium with 2.8μl of 20mg/ml chloramphenicol and let grow for 3.5 hours at 37°C. 2ml of this pre-culture bacterial media was then diluted into 250ml of pre-warmed 2 × Ty media with 175μl 20mg/ml chloramphenicol for 2 hours and 10 minutes and ensured that the OD600 did not exceed 0.6. The culture was cooled down on ice for 5 minutes, divided into four 50ml falcon tubes and centrifuged at a speed of 4000rpm for 5 minutes at 4°C. The cell pellets were suspended in 50ml cold Milli-Q water in each of the four falcon tubes and then centrifuged again at a speed of 4000rpm for 5 minutes at 4°C. After that, the cell pellets were suspended in 50ml cold Milli-Q water in two falcon tubes, and the centrifugation step was repeated as before. A final wash of cell pellets was performed in cold 10% glycerol. After centrifuging for 7 minutes at 4°C and 4000rpm, the supernatant was shaken away and the cells were re-suspended in their own juice.

#### Expressing the mutant library and fluorescence-activated cell sorting (FACS) Sample preparation

0.5μl of 200ng/μl pCIPR plasmids were transformed into 25μl electrocompetent cells made on the same day, inside a 0.1cm-gap cuvette (Bio-Rad) using the Gene Pulser Xcell^TM^ electroporation system (Bio-Rad), with the pre-set protocol for *E.coli* transformation. Cells were recovered in SOC culture media at 37°C for one hour, and an aliquot of the cells was plated on LB agar plate with 100μg/ml ampicillin to examine the transformation efficiency. One transformation with this step produced millions of transformants without creating a bottleneck. Cells were grown overnight in 25ml LB medium with 100μg/ml ampicillin. An aliquot of the overnight culture was diluted 1 in 100 into LB media containing 100μg/ml ampicillin, 0.4% glucose and 0.2% arabinose and grown at 37°C for 2.5 hours to reach an OD600 of about 0.7. The bacteria culture was further diluted 1 in 5 with fresh medium (same composition) and the cells grown for another hour, after which the OD600 was about 0.9. As a control for no CI induction (no repression of the target gene GFP), cells were grown in the LB medium without arabinose but with glucose. All experiments included cells with plasmids containing the wild type CI genotype (positive control) and cells containing an empty pBADM-11 plasmid (to quantify cell autofluorescence) in addition to the cells carrying the mutant library. After the induction of CI expression, cells were immediately diluted 1 in 500 into Phosphate buffered saline (PBS) and put on ice before FACS.

#### Sorting

Sorting was performed at the CRG FACS core facility. A FACSAria II SORP sorter along with the FACSDiva Version 6.1.2 software was used to sort the cells. Bacterial cells were selected based on side scatter (SSC) and forward scatter (FSC), and gate selection was based on FITC-A fluorescence filter for GFP (Figure S9A). Cells were sorted into three gates: the near neutral gate was defined as including 90% of the matching wild type population. The completely detrimental gate included 90% of non-repressed high GFP population (no CI induction). The intermediate population between the two populations mentioned above (about 3~4% of all the library population was in this gate) was also collected (Figure S9B). Purity of sorting was examined by passing the sorted cells through the FACS again immediately after sorting, and recording the population proportions belonging to the sorted gate. At least 30 million cells were sorted per biological replicate.

#### Post-sorting

Sorted cells were kept on ice in PBS in 15ml falcon tube each. They were centrifuged at 4000rpm at 4°C for 30 minutes. The supernatant was removed carefully, and the plasmid-prep was performed directly form the cell pellets. Plasmids from the sorted cells (together with the unsorted input cells) were extracted immediately with the QIAprep Spin Miniprep kit (QIAGEN). The mutagenized region was amplified using barcoded PCR primers (Table S1) for 25 cycles using hot start Phusion polymerase (Thermo Scientific) in 50μl reactions, following the manufacturer instruction. PCR products were purified using the E-gel 2% size-select system (Invitrogen) to remove smaller fragments. In order to produce three full biological replicates, the procedure described up to this -from transformation of the mutation plasmid library to cell sorting and plasmid extraction-was performed three times on three different days (Figure S9C).

Concentration of each purified PCR product was measured on NanoDrop (Thermofisher Scientific). Equimolar quantities of three independent amplifications of the input library (Input) and equimolar quantities of three output replicates from near neutral population (Output1) were pooled together in one Eppendorf tube (Sample1). Equimolar quantities of three output replicates from intermediate population (Output2) were pooled together as a separate sample in a different Eppendorf tube (Sample2). The two samples were sent to EMBL Genomics Core Facility where two PCR-free sequencing libraries were prepared and sequenced on Illumina HiSeq2000 platform. The PCR-free sequencing library Sample1 was run on two lane of an Illumina HiSeq 2500 for each CI concentration experiment. The PCR-free sequencing library Sample2 was multiplexed with other samples to about 10% of one lane loading, considering the small size of the cell population.

#### Verification of mutational effects

22 genotypes (Figure S1C,D and Table S2) were selected based on their enrichment scores and reproducibility (Standard error of the mean enrichment scores <1, see Data analysis section) at both CI concentrations for re-testing. Individual genotypes were constructed using the NEB Q5 site-directed mutagenesis kit (NEB cat. E0554S) with the wild type pCIPR-High and pCIPR-Low plasmids as templates. After verifying the sequences by Sanger sequencing, we picked four colonies from each genotype to examine their target gene GFP expression levels (Figure S1C, D). The experiment was performed in one batch on the same day so that the results from this experiment could be used as a confirmation set to which other FACS experiment sets can be mapped. LSR Fortesta florescence analyser was used at the CRG FACS Core facility.

GFP signal and the shape information of 10,000 cells per biological replicate were recorded, and the “.FCS” files from the recordings were analyzed using the *FlowCore* package in R. Cells were filtered based on SSC and FSC, and the first 3,000 cell recordings were discarded to avoid cross-well contamination. The mean output GFP signal (in AU, arbitrary units) from about 5,000 cells in each biological replicate of individual variant was calculated after the filtering process. The mean GFP signal and standard error of the mean for each variant were obtained from each biological replicate.

#### Quantification of CI protein expression

The relative amount of CI protein at the two expression levels was quantified by tagging CI with GFP at its C-terminus with the flexible linker amino acid sequence GSAGSAAGSGEF (Waldo et al., 1999). The PR-GFP sequences were removed from the original pCIPR-High and pCIPR-Low plasmids to make plasmids pCIGFP-High and pCIGFP-low. Fluorescence from CI-induced cells was analysed using a LSR Fortesta florescence analyser at the CRG FACS Core facility (Figure S10A). In the same experiment, GFP calibration beads (CloneTech) were used to calibrate and obtain exact molecule numbers based on the GFP signal (Figure S10B, C). For quantification, mean GFP signals and standard errors of were calculated from four biological replicates.

### Data analysis

#### From sequencing data to target gene expression

Our data analysis pipeline consists of three main parts: 1) Filtering. 2) Mapping enrichment scores to the target gene (GFP) expression levels. 3) Correcting for the batch effects (Figure S9C) and the detection limits set by the experiment. The processed final datasets for the analysis were organised both on nucleotide level and amino acid level. Even though our conclusions were mainly based on amino-acid level mutational effects, the dataset with nucleotide-level mutational effects was needed as reference.

The analyses from sequencing data to GFP expression level were all performed on the nucleotide-level, and the amino-acid level mutational effects were examined based on the processed nucleotide-level datasets. Whenever involving combining replicates (at the level of enrichment scores, predicted GFP singles at the nucleotide level and at amino acid level), the random error model was used.

#### From Illumina sequencing reads to variant counts

To extract variant counts from the raw sequencing data, we adapted the pipeline developed by our group in a previously published project (Julien et al., 2016). Specifically, the raw sequencing data was demultiplexed with the SABRE software (https://github.com/najoshi/sabre) and paired reads were merged with the PEAR software (Zhang et al., 2014) with parameters set not to allow any mismatches in the overlap regions. Reverse complementation of merged sequences was performed when necessary with the fastx_reverse_complement tool (http://hannonlab.cshl.edu/fastx_toolkit/). Then the primer sequences were trimmed using the seqtk tool (https://github.com/lh3/seqtk). Finally, the number of occurrences of each variant was counted with fastx_collapser (http://hannonlab.cshl.edu/fastx_toolkit/) and a custom python script(Julien et al., 2016).

#### Calculating enrichment scores and filtering

Variants up to 2-Hamming-distance nucleotide changes from the wild type sequence with at least 100 read counts in all three input replicates were selected for further analysis (Figure S11A,B). The 100 read count threshold included all the 1-Hamming-distance nucleotide changes (n=531) but only about 11% (n=10,862 for low expression dataset) and 7% (n=3,686 for high expression dataset) of all the 2-Hamming-distance nucleotide changes observed. This restriction was necessary to obtain the confident variant counts. The threshold was chosen based on the logic that each bacterial cell be expected to carry hundreds of plasmid copies (pUC replication origin). Considering experimental steps of plasmid extraction and PCR amplifications until obtaining read counts from Illumina sequencing, we reasoned that variants observed less than 100 read counts were likely to be from too few cells, resulting in unreliable enrichment scores for the following steps.

Enrichment scores for each variant *v* from each experimental replicate *i* (REPi), for each sorted cell output *j* (*Oj* with *O1* as near neutral fraction and *O2* as partially detrimental fraction) were calculated as follows:

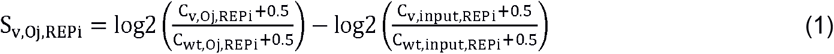

With *C* as sequencing read counts, *v* as variant, *wt* as wild type. A pseudo count of 0.5 was added to avoid log 0. Poisson-based error for each variant for each replicate for each output (*SE*_*v,Oj,REPi*_) was also calculated using the formula below:

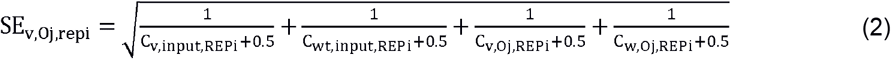

In order to merge scores over replicates for each output and for each variant, and to be able to filter variants based on the standard errors of the mean, a random-effect error model as proposed by Rubin *et* al for this type of data analysis (Rubin et al., 2017) was used.

The details are as follows:

For the first iteration, for each output, an initial error 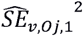 for each variant was calculated based on its standard deviation from the unweighted mean.

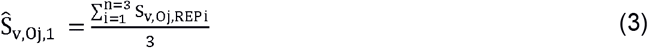

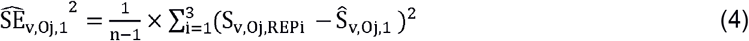

The initial weighted mean enrichment score for each output was calculated as the follows:

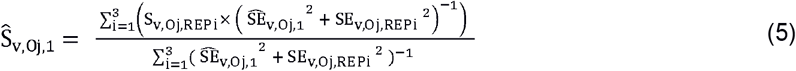

For each iteration *k*, the standard error was calculated as follows:

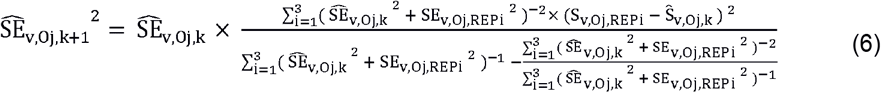

After 50 iterations (*k*=50), the final mean enrichment score and standard error for each variant for each output were calculated as shown in equations (7) and (8) respectively.

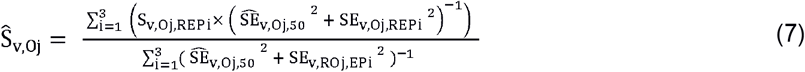

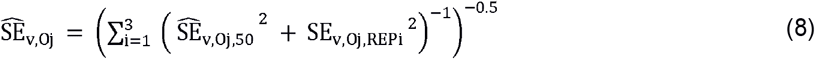

In order to estimate the overall errors of enrichment scores for each variant and to filter only the confident data for the following data analysis, the estimated errors from Output1 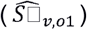 and Output2 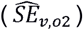 were combined with the following formula:

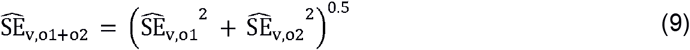

Variants with 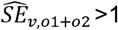 were removed for downstream analyses (Figure S11C, D).

#### Mapping enrichment scores to GFP signal

In order to calculate GFP signals from enrichment scores, we first examined the relationships between GFP signals and enrichment scores from individually assayed confirmation data set (Table S2). As designed by the experiment, the smaller enrichment score from the Output1 S_v,o1_ was, the higher GFP signal (more detrimental) of a variant was (Figure S12A). Enrichment scores from the Output2 S_v,o2_ (the intermediate fraction) did not relate monotonically to the mean GFP signal, because variants enriched in Output2 (S_v,o2_) were depleted for both strongly detrimental and near neutral variants (Figure S12B).

To examine the possibility of predicting GFP signals with a linear combination of the two enrichment scores for each replicate from each expression level experiment, we built linear models to predict the mean GFP signals with S_v,o1,REPi_ and S_v,o2,REPi_ with the confirmation dataset. The calculated GFP signal from the mean enrichment scores predicted the individual variants’ GFP signals well (Figure S12F). However, the predictions were not completely linearly related with the observed GFP signals.

In order to improve the GFP signal predictions based on the enrichment scores, for each biological replicate, we transformed each S_v,o2,REPi_ to S_v,o2,trans, REPi_ based on its relationship with S_v,o1,REPi_ such that variants predicted to be detrimental by S_v,o1,REPi_ would have higher S_v,o2,trans, REPi_ and variants predicted to be near neutral by S_v,o1,REPi_ would have lower S_v,o2,trans, REPi_ (Figure S12D, E).

The logic behind this transformation was as follows: 1) A potentially beneficial mutation was expected to be enriched in Output1 and depleted in Output2 (S_v,o1,REPi_ > 0 and S_v,o2,REPi_ < 0). We kept the Output2 score as it was. 2) An intermediately detrimental mutation was expected to be enriched in Output2 (S_v,o2,REPi_ > 0) regardless of its enrichment score in Output1. We kept its enrichment score in Output2 as it was as well. 3) A very detrimental mutation was expected to be depleted both in Output1 and Output2 (S_v,o1,REPi_ < 0 and S_v,o2,REPi_ < 0). In order to distinguish S_v,o2,REPi_ of these variants from that of potentially beneficial mutations (the first case, where S_v,o2,REPi_ is also smaller than 0), we transformed S_v,o2,REPi_ to a positive value and bigger than the intermediately detrimental variants’ S_v,o2,REPi_. This way, a transformed S_v,o2,trans, REPi_ was expected to be bigger for more detrimental mutations (Figure S12C). To avoid influence by extreme outliers, 95^th^ quartiles (Q) were used as thresholds for detrimental mutations (S_v,o1,REPi_ < Q(S_wt_syn,o1,REPi_, 0.95)) and as an approximate for the maximum S_v,o2,REPi_ before transformation. To summarize, the equation follows:

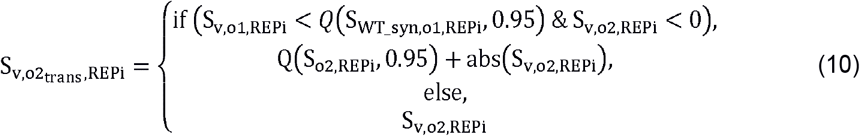

A linear model was built again to predict the mean GFP signals for each expression level experiment with the mean enrichment scores 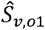 and 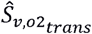 using the confirmation dataset. Inverse of the variance was used as weights. This linear model improved the prediction of GFP signal in the low CI expression dataset (Figure S12G). For the high expression dataset, the _v,02trans_ coefficient was not significant (Table S3, Figure S12G) and including the S_v,o2_trans,REPi_ did not improve logged GFP signal (as an output of mutational effects, denoted *O*) O_v,REPi_ predictions (note R^2^ and the median RMSD did not change in the predictions for high expression dataset, Figure S12G). Therefore, we set S_v,o2_trans,REPi_ = 0 when calculating signals and the errors for the high CI expression dataset in the following equations to avoid inflating the errors of the estimation (refer to equation (12)).

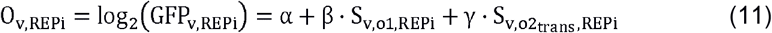

*O_v,REPi_* above is the output GFP signal in log scale for each variant in each of the three biological replicates *i* and the coefficients α, β, γ (Table S3) derived from the linear model trained with the confirmation dataset.

A measurement error for the log GFP signal (*OE_v,REPi_*) for each variant *v* in each replicate *i* and for each CI concentration (high and low) was calculated with the following formula:

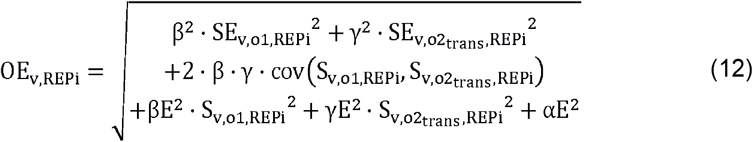

Where βE^2^, γE^2^, αE^2^ are squares of the standard errors of the estimated α, β and γ coefficients respectively, and cov(S_v,o1,REPi_, S_v,o2,trans,REPi_) is the covariance between S_v,o1_ and S_v,o2,trans_ for each replicate (Table S4).

#### Correcting technical biases

Each biological replicate from FACS sorting on different days had different ranges of GFP expression levels (GFP index, Figure S9C) and these biases were reflected on the estimated *O_v,REPi_* (Figure S13A,B). In order to correct these technical biases, one replicate from each CI concentration experiment was set as reference, and the other replicates were linearly mapped to the same range as the reference replicate (i.e., replicate 2 as the reference).

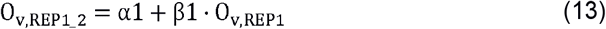

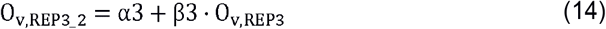

In the function above, the coefficients 1 and 1 derived from mapping the line defined by replicate 1 replicate 2 wild type _wt,REP2_and weighted means of the nonsense mutations’ _n0n,REP3_ (Table S5, Figure S13A - C).

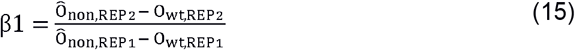

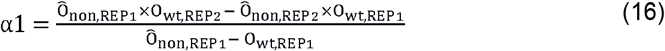

The same equations as (15) and (16) applies to coefficients α3 and β3 to map replicate 3 to replicate 2 by only substituting replicate 1 with replicate 3.

The mean GFP signals 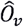 and standard errors of the mean 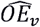 over biological replicates were calculated using random-effect error model as described in the previous section for combining enrichment scores over the biological replicates.

### Calculating mutational effects at amino acid level

In order to examine mutational effects at the amino acid level, the processed data at the nucleotide level was converted to the amino acid level.

First, for each replicate, weighted mean GFP signals of all the nucleotide variants encoding the same amino acid variants were calculated. The inverse of the GFP signal errors of the nucleotide variants were given as weights. Errors from each nucleotide variants were propagated as the error of the GFP signals for each amino acid variant in each replicate.

Then, mean GFP signals 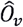 and the standard errors of the mean 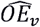 over biological replicates at amino acid level were calculated based on the random-effect error model as for combining enrichment scores and nucleotide level GFP signals over replicates.

### Rescaling the mean GFP signals to the detection limits

In the FACS experiments, the detection limit for the lowest GFP signal was equal to the auto-fluorescence of the bacterial cells not expressing GFP. The auto-fluorescence of the bacterial cells was not distinguishable from the cells that repressed the target gene GFP expression completely (CI WT high expression) (Figure S9C). The theoretical maximum GFP expression level was equal to that of bacterial cells expressing the target GFP without any repressor.

However, some variants’ estimated GFP signals from the bulk sequencing data exceeded the GFP signal range defined by theoretical maximum and minimum GFP. These GFP signals outside the theoretical limits were not likely to be real and they could potentially bias our analysis.

In order to correct this problem, estimated GFP signals from the enrichment scores were rescaled to abide to the theoretical maximum and minimum GFP ranges. The lower GFP detection limit was determined by the lower limit of 95% confidence interval from the mean CI WT high expression GFP level. The upper GFP detection limit was determined by the 95^th^ percentile of the weighted mean GFP signals of all nonsense mutations at low expression level of CI. The 95^th^ percentile (or confidence interval) rather than the mean WT or nonsense GFP signals were selected as detection limits, so that the modes of the mutational effects would not shift after rescaling.

This GFP detection range [4.5,12.8] was first divided into 1000 evenly spaced bins (*O_k_*). Then, given the observed mean GFP signal and the standard error of a variant, the probability of the true mean GFP signal of the variant falling into each bin was calculated as follows:

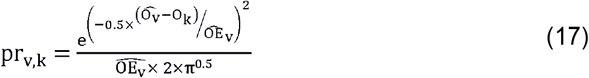

Finally, the mean GFP signal of a variant was calculated based on the weighted mean of the GFP signals from each bin with the weights given as the probability of the true mean falling into each bin *k* (pr_v,k_), as shown below:

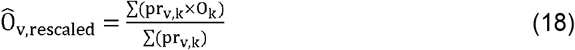

The 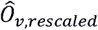 (Figure S13D) was used as the mean GFP signal for each variant in the following analysis, denoted as 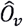 replacing the value before transformation, and the standard error 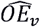 was kept the same as before rescaling.

### Folding energy, binding energy and structural analysis

Folding energy prediction and structural analysis were performed based on the 3.909Å x-ray structure (PDB 3BDN) of CI dimer bound to an operator site OL1.

To estimate the mutational effects on folding energies and binding energies of CI protein, we used FoldX4 software (Schymkowitz et al., 2005). First, *BuildModel* command was used to build a structural model from each single mutation in our experiment. Then, the *AnalyzeComplex* command (with the *complexWithDNA* option set to *true*) was used to obtain the absolute energies of protein-DNA complex (ΔG_CI-OR,FoldX_) as well as the protein itself (ΔG_F,FoldX_) for each mutation. Binding energy of CI to DNA (ΔG_B,FoldX_) was calculated as energy difference between the protein-DNA complex and the protein by itself for each mutation. ΔΔG for folding (ΔG_F,FoldX_) and binding energies (ΔG_B,FoldX_) for each variant were calculated by subtracting folding and binding energies of wild type CI respectively.

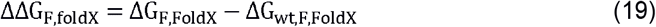

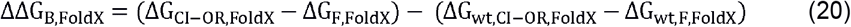

Analyses were repeated with PDB structure 1LMB (1 Å x-ray structure of CI N-terminal domain bound to OL1) and with 3BDN structure bound to OR1 instead of OL1 (by mutating OL1 sequence to OR1 based on PDB 3BDN structure). FoldX4 returned the same ΔΔG with these analyses; therefore, only results using PDB 3BDN as a template were shown.

3D structures were visualized and analysed using PyMOL (v1.7.6.0). Amino acid positions were classified as core residues if the ratio between solvent exposed area and the total area fell within the first quartile of the obtained data based on a PyMOL script (“get_area”, https://pymolwiki.org/index.php/Get_Area) with parameters dot_density set as 4 and dot_solvent set as 1. Positions were classified as DNA-contacting when the differences in the solvent exposed area without DNA and with DNA were greater than 0.1Å^**2**^.

### Other features tested for their association with mutational effects

562 amino acid indices taken from the AAindex database (https://www.genome.jp/aaindex/) (Kawashima et al., 2007) together with BLOSUM62 matrix scores (ftp://ftp.ncbi.nih.gov/blast/matrices/), structural information, and FoldX predicted energy values were examined. The top features that correlated with the mutational effects of CI protein were: (1) the hydrophobicity index (Zaslavsky et al., 1982); (2) the number of negative charges introduced by a mutation (Cherstvy, 2009); (3) the amino acid substitution matrix BLOSUM62 (Henikoff and Henikoff, 1992); (4) changes in the protein folding energy; (5) changes in the protein-DNA binding energy predicted by FoldX (Schymkowitz et al., 2005) together with the structural features of mutations (i.e., at the core, interface with DNA or at the solvent-exposed positions).

### Mathematical model

Our aim was to build a mathematical model that captures the most important features of the system that apply to all mutations. The model propagates the effects of mutations on the folding of the lambda repressor to changes in expression of the target gene through the well-described regulatory model of the PR promoter. The model makes the following assumptions: 1) Mutations change the free energy of protein folding so altering the fraction of folded protein; 2) the fraction of folded protein is independent of the protein concentration; 3) changes in protein folding free energy are additive for all mutations. In reality, all of these assumptions may be violated for some mutations. For example, some mutations will also affect the binding affinity of the lambda repressor to the DNA operator sites or alter transcription or translation. Others may result in protein aggregation. Moreover, the fraction of folded protein may not be independent of concentration, for example at very high expression levels because of chaperone titration. Finally combining mutations in structurally contacting or indirectly energetically-coupled residues may result in non-additive changes in free energy. However, our aim was to test whether the simplest possible model of the system captured the overall changes in mutation effects and changes in the strength and sign of genetic interactions as the expression level changed. We of course acknowledge that some mutations will not meet these assumptions and these exceptions likely contribute to some of the unexplained variance in our data.

### Regulatory interaction model of the CI-repressor system

Ackers’ 8-configuration model (Ackers et al., 1982) was used to predict the relationship between the total amount of CI protein and the expression levels of its repressed gene. As in our experiment, the CI regulatory interaction system in Ackers’ model involves three operators (OR1, OR2 and OR3), resulting in eight possible configuration states (CS) in which the CI dimer can bind to the operators (Table S6). Based on the model, each configuration state causes the downstream promoter to be in either an ON or OFF state (Figure S3A). Only two configuration states fail to repress expression of the target gene: when the CI dimer is not bound to any operators (CS1) and when CI dimer is only bound to OR3 (CS2). The probability of repressing the target gene expression is the sum of the probabilities of the six remaining configuration states that result in the OFF state of the promoter. The likelihood of each configuration state is a function of the binding energies and the free CI protein dimer concentration when the number of OR binding sites is fixed. In Ackers’ model, the number of OR sites is equivalent to that found in an average lysogen (bacteria that carries the phage genes integrated in its genome) with the ORs integrated into its genome (Ackers et al., 1982).

The probability that each of the eight configuration state (*f*_*CSi*_) to occur is:

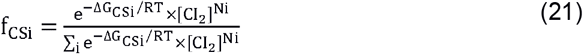

Where Δ*G*_csi_ is the total free energy of lambda repressor dimers in the respective configuration *i*; the exponent *Ni* is the total number of the lambda repressor dimers in the corresponding configuration *i*; [CI_2_] is the free dimer concentration; R is the gas constant (R = 1.98×10^-3^ kcal/M) and T is the absolute temperature (310.15 kelvin).

The probability of repression (P_s_) is the sum of the probabilities of the configurations in which promoter P_R_ is repressed (∑_i={3:8}_*f*_*CSi*_). To calculate P_s_ as a function of the free dimer CI concentration [CI_2_] based on the equation (19) and Table S6, we obtain the following equation:

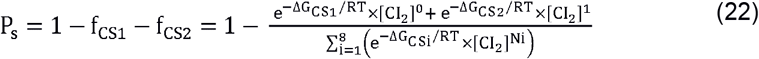

The target gene expression level (GFP) is modelled to be proportional to the binding probability of the RNA polymerase, which is given by one minus the probability of repression by CI (P_s_) (equation 21).

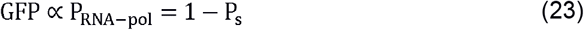

Despite its simplicity, this model has been shown to be predictive of the gene expression levels (Bintu et al., 2005). Because bacterial cells displayed auto-fluorescence (GFP_auto_), this auto-fluorescence signal from bacteria needed to be considered when measuring the effects of mutations on GFP levels. Therefore, by rewriting the equation (21) by taking into account the auto-fluorescence of the cells, the probability of GFP repression can be shown as in the equation (22).

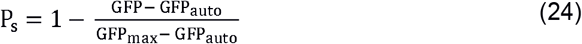

Both equations (20) and (22) show the probability of repressing the target gene, with equation (22) as a function of the GFP signal and equation (20) as a function of the free CI dimer concentration. By combining equation (20) with equation (22), we obtain an equation that describes the relationship between the free CI dimer concentration [CI_2_] and the GFP signal as shown in the following equation:

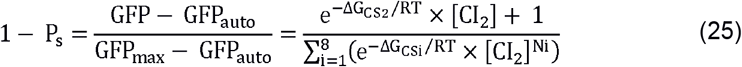

By rewriting the equation (23), we can show the GFP signal as a function of free CI dimer concentration:

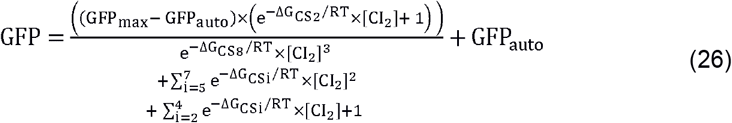

The equation (24) allows us to calculate the GFP signal for each variant from a known free CI dimer concentration. In order to calculate [CI_2_] from equation (24*), uniroot* function was used with R script to find a unique root of equation (24) that was within the range of 10^-40^ and 10^-3^ (M).

Next, the relationship between the total CI concentration [CI_T_] and the free dimer concentration [CI_2_] was evaluated, in order to model the relationship between the GFP signal and total CI concentration. This is because the total protein concentration in the cells but not the free dimer concentration of the protein is the one that can be experimentally measured and manipulated. The total lambda repressor concentration in the cell [CI_T_] is the sum of the free monomer concentration [CI] plus two times the concentrations of the free dimer [CI_2_] plus two times the concentration of the dimers bound to operators [OR]. Compared to the original Ackers’ model, in our experimental system, each bacterial cell was expected to carry up to hundreds of folds more operator sites, the same fold changes in CI protein coding region and the target gene. Given the same fold changes in all the functional blocks in this model, we simply kept the same parameters from original model and mapped our experimental system to the original model system.

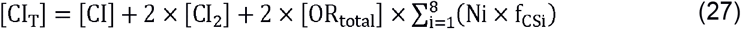

The concentrations of free monomer [CI] and free dimer [CI_2_] follow the equilibrium:

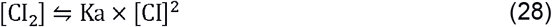

By combining the equations (25) and (26), we can describe the relationships between [CI_T_] and [CI_2_] as follows:

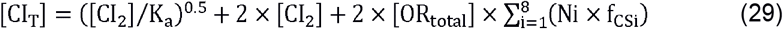

By further substituting 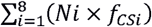 from the equation (27) with the equation (19), we obtain the following equation:

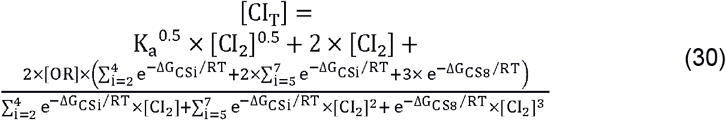

The equation allows us to calculate [CI_T_] from [CI_2_]. [CI_2_] can be calculated by finding the unique root from the equation (24) from the known GFP signal as described earlier. Given the complexities of both equations (24) and (28), the calculations were performed in two steps according to the two equations. For the following process, for ease of reference, we denote the process of calculating total protein [CI_T_] for each variant from its target GFP signals *f’_Ackers_*:

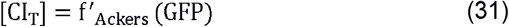

The reverse process to calculate GFP signals from the total protein [CI_T_] involves two steps: 1) inversing equation (28) to calculate the corresponding [CI_2_]; 2) calculating GFP with equation (24) with [CI_2_] from the previous step.

Inversing and finding the exact root of equation (28) is mathematically impossible. Therefore, an approximate solution was found based on a local polynomial regression (*loess* function with R, span parameter 0.3) describing the relationship between [CI_2_] and [CI_T_] based on equation (28) (Figure S14A).

Based on equation (24), GFP signal was calculated by inputting [CI_2_] from the previous step. We denote the process as *f_Ackers_*, which is the inverse of equation (29) for the ease of future reference.

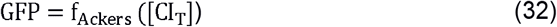

The parameters were kept as they were originally used in the model by Ackers (Ackers et al., 1982) (Table S7) that were experimentally determined.

Two additional parameters (GFP_max_ and GFP_auto_) were specific to our experiment and not described in the original model by Ackers. For modelling, the maximum GFP signal GFP_max_ was defined as the weighted mean GFP signals of all single nonsense mutations with weights given as the inverse of the variance (3470.67 AU, or 11.76AU in log2 scale) based on the CI low expression dataset. The minimum GFP signal GFP_auto_, corresponding to the cellular auto-fluorescence GFP signal, was found through parameter search as follows. To start with, two constraints for GFP_auto_ were considered: first, based on the regulatory interaction model, repression of the target gene expression can never reach 100% even though it can infinitely approach this level.

In other words, the GFP_auto_ cannot be set to be the same as the GFP signal from the wild type protein at high expression. Second, GFP_auto_ should allow the calculated ratio of wild type [CI_T_] between high and low expression levels based on equation (29) *f’_Ackers_* to agree with experimentally quantified ratio (15:1, see protein quantification section, Figure S10). We performed the parameter search for GFP_auto_ that allowed the ratio of calculated wild type [CI_T_] at two expression levels to be 15:1 based on the model calculation as shown below:

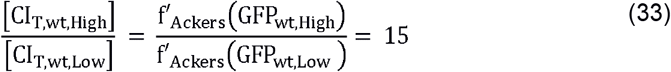

GFP_auto_ was estimated to be 23.24AU (4.54AU in log2 scale) to meet the condition set by equation (31).

### Estimating the functional protein concentration for all variants

An estimate of wild type CI protein concentration [CI_T,wt_] in each of the two experiments can be obtained by inputting GFP_wt,High_ and GFP_wt,Low_ values into ***f’***_Ackers_ function. The same way, the total protein concentration of a variant [CI_T,v_] can be derived for each experiment with the ***f’***_Ackers_ function. Differences between [CI_T,v_] and [CI_T,wt_] were assigned to differences in their functional protein fraction rather than changes in the total expressed protein amount. This is based on the assumption that the mutations with one or two amino acid alterations affected GFP levels mostly through changing the fractions of natively folded protein (*f_N_*).

In order to calculate the fraction of correctly folded protein for each variant (*f_N,v_*), knowledge of the total expressed protein concentration [CI_E_] for each experiment was needed. Based on the calculated [CI_T,wt_] at both low and high expression levels and the information that the fraction folded of the wild type protein is 0.9913 (see the next section), the total expressed protein concentration in the cell can be calculated by multiplying the concentration of functional wild type CI protein by 1/0.9913 (Table S8).

The fraction of natively folded protein for a variant *v* (*f_N,v_*) was calculated as the ratio of [CI_T,v_] (that is calculated based on ***f’***_Ackers_ (GFP_v_)) over total expressed CI [CI_E_] (as a parameter calculated based on ***f’***_Ackers_ (GFP_wt_), Table S8):

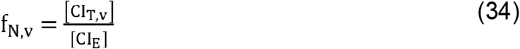

### Thermodynamics of CI folding model

CI has been shown to follow a two-state model of protein folding(Huang and Oas, 1995) that can be described with the following equation:

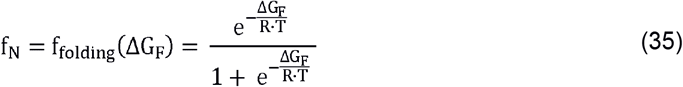

With f_N_ as the fraction of natively folded protein, ΔG_F_ as the total free energy of the protein folding. R is the gas constant (R = 1.98×10^-3^ kcal/M) and T is the absolute temperature of our experimental setting (T=310.15 kelvin, 37°C).

Rewriting the equation (33), we obtain:

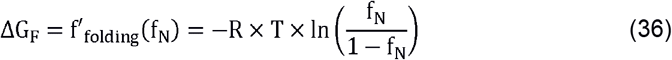

The equilibrium between the concentration of unfolded and native CI protein follows the equation below:

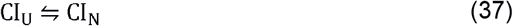

Equation (35) is governed by an equilibrium constant K_fold_ whose value is known to be 114 for the wild type CI protein(Parsell and Sauer, 1989) :

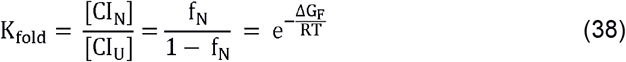

By solving equation (36) with K_fold_ = 114, we obtain the wild type CI f_N_ = 0.9913 which was used to calculate the total protein concentration in the cells (Table S8), as shown in the previous section.

The folding energy of a double missense mutation (AB) can be predicted by adding the folding energies of the two single mutations (A and B) that together make the double mutation (AB).

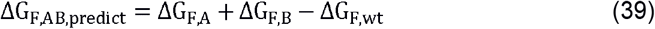

### Combining thermodynamics of the CI folding model with the regulatory interaction model of CI-OR system

To predict GFP signal of a mutation *A* from ΔG_F_ values of the mutation, the output of ***f***_folding_ function was added to ***f***_Ackers_ function:

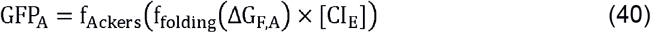

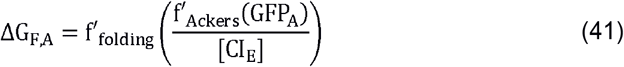

### Comparing four different sub-models for the effects of mutations

To evaluate the importance of (a) protein folding and (b) CI-concentration-dependent repression of the target gene expression independently as well as in combination, we generated and compared four models (Figures 3, S3). The four models are based on four different assumptions. The first model is the log-additive model where changes in the target gene expression levels are simply additive in the log scale (Figure S3C). The second model is the full model that incorporates the effects of mutations both at the level of protein folding and at the level of regulatory interaction of CI-OR system on the target gene expression (as shown in equations (37–39), Figures 3A, S3B,C). The third model is a protein folding-only model that incorporates the thermodynamics of protein folding but not the regulatory interaction model (it assumes a linear relationship between target gene expression and functional CI concentration). Therefore, the protein folding energies are additive features of this model (Figure S3B-D). The last model is the regulation-only model that incorporates the regulatory interaction model but not the thermodynamics of protein folding (it assumes a linear relationship ΔG_F_ and f_N_). Therefore, the functional protein amount is the additive feature of this model (Figure S3 B,C,E).

Depending on the model evaluated, the functions linking the target gene GFP expression level to [CI_T_], or [CI_T_] to ΔG_F_ can be different. The details of each model are explained below.

#### 1) Log-additive model

Consistent with extensively used null models where the effects of mutations are log-additive, this model predicts the log GFP signal of a double mutation *AB* relative to the wild type to be the sum of the log GFP signals of each of the two single mutations relative to the wild type:

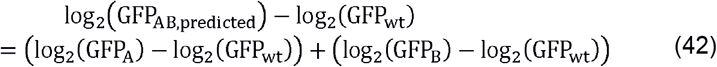

Therefore,

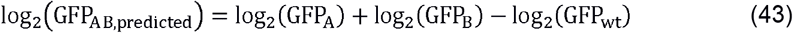

#### 2) Full model

To predict the GFP expression levels of a double mutation, we first estimated the ΔG_F_ of the corresponding single mutations using equation (39). The ΔG_F,AB_ of the double mutation was calculated with equation (37) and then converted to an expected GFP signal using equation (38).

#### 3) Folding-only model

This model assumes that the GFP expression levels are linearly responsive to the fraction of natively folded protein f_N_. That is, this model replaces ***f***_Ackers_ with a linear transformation between GFP signal and f_N_ (Figure S3E). At the same time, this model includes the nonlinear relationship between f_N_ and ΔG_F_ that was introduced by the thermodynamics model of protein folding. Thus, for a mutation *A*, the relationship between GFP signal and the fraction of folded protein f_N_ was given by a modified version of ***f***_Ackers_, which we call ***f***_model3_.

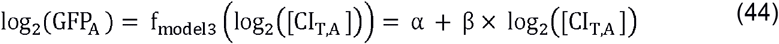

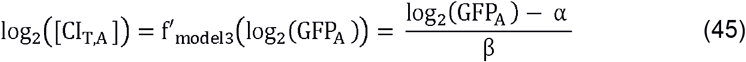

The output of ***f***_model3_ can then be introduced into ***f’***_folding_ (equations 34 and 37).

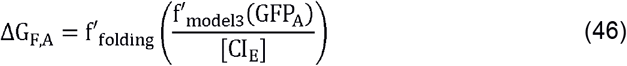

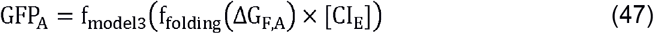

The α and β parameters from ***f***_model3_ (equation (42)) determine the linear relationship between the functional repressor concentration and GFP expression levels (α is the intercept and β is the slope). Also, the parameters [CI_E,low_] and [CI_E,high_] (Table S8) were kept the same as in the other models.

Comparing the mutational effects at two expression levels based on equation (42), we obtain the following equation:

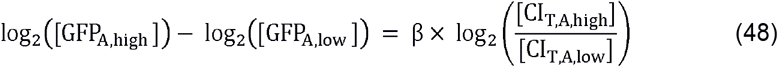

The ratio of [CI_T,A_] at two expression levels was set as the constant 15 (as defined by wild type protein, see the previous section). Equation (46) therefore can be re-written as follows:

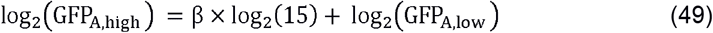

By substituting β×log_2_(15) with a coefficient *C*, we can rewrite equation (47) as follows:

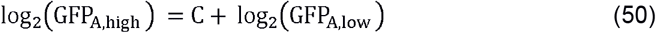

From equation (48), we can see that GFP signal at the two CI expression levels is linearly related with the fixed slope of one in the log space. Parameter search was performed to find the coefficient *C* that best described the observed relationships between GFP signals at low and high expression levels of CI. In detail, we firstly sampled a hundred log_2_(GFP_v,low_) values ranging between log_2_(GFP_wt,low_) = 7.23 and log2(GFP_max_) = 11.76. Then, a range of intercept *C* between −3.3 and −1.3 with the step of 0.03 was used to calculate the corresponding log_2_(GFP_high_) for each log_2_(GFP_low_) (Figure S14B-D). The value *C* =-2.07 was selected that resulted in the smallest sum of the squared distances from the observed data points to the line defined by simulated relationship between GFP signals at two expression levels (Figure S14C). Based on *C*=β×log_2_(15), we further calculated β = −0.52.

The coefficient α was calculated by placing the values β and the wild type CI low expression data [CI_E,low_] and log_2_(GFP_wt,low_) to equation (42) and rewritten as below:

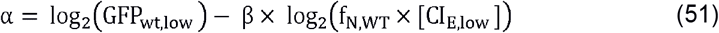

With *log*_2_(*GFP*_*wt,l0w*_ as observed in the experiment, β = −0.52 as calculated above, and the known parameters f_N,wt_ = 0.9913 and [CI_E,low_] = 5.5×10^-8^, we obtained α = −17.

To estimate the GFP signals of a double mutations with the folding-only model, we first estimated the ΔG_F,A_ and ΔG_F,B_ of the corresponding single mutations using equation (44). Double mutants’ ΔG_F,AB_ was calculated using equation (37). ΔG_F,AB_ of the double mutant was then converted to an expected GFP signal using equation (45) (Figure S3B,C).

#### 4) Regulation-only model

This model assumes that the f_N_ of a protein is linearly related to its ΔG_F_. That is, this model replaces ***f***_folding_ with a linear transformation between f_N_ and ΔG_F_. At the same time, this model includes the nonlinear relationship between f_N_ and GFP expression levels from Ackers’ model.

Because of the assumed linear relationship between f_N_ and ΔG_F_, the effects of mutations are additive in f_N_ space making ***f***_folding_ unnecessary in this model at all (Figure S3B,C). To estimate the GFP expression levels of a double mutant with the regulation-only model, we first estimated the functional protein concentration of the corresponding single mutants using ***f’***_Ackers_ (GFP). The expected functional protein concentration of the double mutant was then given by the following equation.

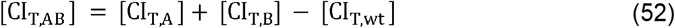

The expected GFP signal for this double mutant was calculated using ***f***_Ackers_ ([CI_T,AB_]), as shown in equation (30) (Figure S3B, C).

### Simulating mutational effects and genetic interactions based on the model

To test to which extent each model can explain 1) the double mutational effects given the single mutational effects 2) the relationship between the mutational effects at the two protein concentrations 3) the pair-wise genetic interactions at both protein concentrations, we simulated mutational effects and their interactions based on each model to compare with our data.

### Simulating the mutational effects based on the model

We sampled 100 ΔG_F_ values equally spaced between −3kcal/mol and 3kcal/mol, and estimated their GFP signals at high and low CI concentrations using each of the four sub-models described above. For a given model, plotting the GFP signals predicted for the high CI concentration case against the GFP signals predicted for the low CI concentration case resulted in a curve (or a line) (Figure 3E).

To test how well each model explained the observed protein concentration dependent mutational effects, we used the *Princurve* package(Hastie and Stuetzle, 1989) in R to calculate the sum of squared distance from the curve (SSDC) between every experimental data point and the line or curve described by the model (Figure S14E).

To predict the ΔG_F_ for a specific variant, we first projected each data point in the log GFP (high CI concentration) vs. log GFP (low CI concentration) scatter plot to the nearest point in the model curve (line, in the case of folding-only model) (Figure S14D, E). The projected GFP signal corresponds to a single ΔG_F_ of each variant based on the model. This correction allowed us to estimate a single ΔG_F_ using the GFP value for both the high and the low CI concentrations. Finally, this estimated ΔG_F_ (functional protein concentration [CI_T_], in the case of regulation-only model) of a variant was used in the following processes for predicting double mutational effects and to predict the epistasis patterns.

### Comparing model predicted and observed double mutational effects

The percentage of variance explained (PVE) for the mean GFP expression levels of the double mutation was calculated as follows:

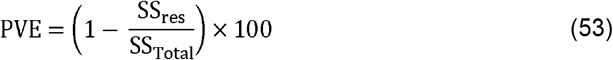

Where *SS_res_* is the residual sum of squares between the model-predicted versus the observed GFP expression levels and *SS_Total_* is the variance in the observed data.

### Predicting pair-wise genetic interactions with each sub-model

Epistasis was defined as the difference between the GFP expression levels of a double mutant based on the model (full model, folding-only model and regulation-only model) and the log-additive model (equation (41)), as shown in the equation below (Figure 2F).

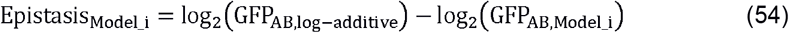

For a given double mutant, we first predicted the ΔG_F_ values (full model and the folding-only model) or f_N_ (regulation-only model) of the corresponding single mutants, as stated above. We then used each model to convert the double mutant’s predicted ΔG_F_ or f_N_ value back into the GFP signal. This predicted GFP signal was compared with the expected GFP signal based on the log-additive null model (equation (41)). The genetic interaction patterns were further compared to the experimental observation (Figures 3G, H, J, K and S4, S5).

The summary of the modelling mutational effects based on each model was illustrated as a cartoon in the Figure S3A–C.

### Toy models of three protein expression–fitness relationships

Three most common fitness-protein concentration relationships were modelled based on the fitness effects of changes in protein concentrations in yeast (Keren et al., 2016).

Fitness increases with lower protein concentration:

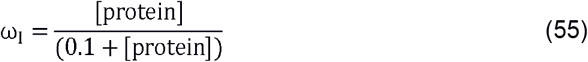

Fitness with optimal protein concentration:

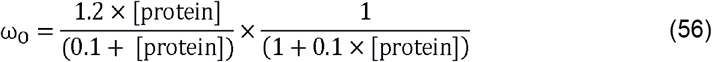

Fitness decreases with higher protein concentration:

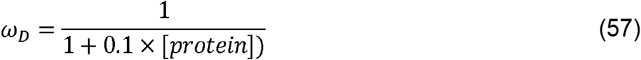

These functions were integrated into the full model (sub-model 2) in place of ***f***_Ackers_ to build three new models linking fitness to changes in protein folding energy ΔG_F_. Note that because ***f***_folding_ was left untouched, all these three new models assumed a two-state protein folding kinetics as for CI protein. Mutational effects and pairwise genetic interactions were analysed at two simulated protein concentrations (high and low) based on these models. For both ‘Increasing’ and ‘Decreasing’ fitness landscapes, the two wild type protein concentrations in each simulation were selected so that one wild type protein concentration would be abundant enough to be robust to mutational effects and the other one would be sensitive to the mutational effects. For the ‘Peaked fitness landscape’, the two protein concentrations were selected so that the fitness effects would be the same but the protein expression levels at ‘Low’ would be below the optimal protein concentration and at ‘High’ would be above the optimal protein concentration.

We evaluated the effects of 50 mutations with ΔΔG_F_ evenly spaced between −1kcal/mol and +5kcal/mol in four different wild type proteins with different protein folding energies: (1) very stable wild type protein (ΔG_F,wt_ = −3kcal/mol); (2) stable wild type protein (ΔG_F,wt_ = −1.6kcal/mol); (3) marginally stable wild type protein (ΔG_F,wt_ = −1kcal/mol); (4) unstable wild type protein (ΔG_F,wt_ = 0kcal/mol) (Figures 5A – H, S7). The effects on fitness of all the pairwise combinations of mutations were also evaluated assuming that the effects of mutations are additive in ΔΔG_F_ space.

Epistasis was quantified as the difference between the “observed” double mutational effects (calculated by adding the ΔΔG_F_ of the single mutations, as described above) with the expected effects (calculated by adding up single mutational effects based on the log-additive null model):

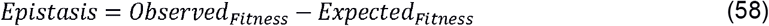

## QUANTIFICATION AND STATISTICAL ANALYSIS

Statistical details of experiments including the statistical test used, the exact number of the data points, mean values, standard errors of the mean (SEM), and 95% confidence intervals, p-values can be found in the figure legends and results. As described in the Data Analysis section of the STAR Methods, data with low reproducibility from the three biological replicates (SEM>1 for the predicted mean GFP) were excluded from subsequent analyses.

## DATA AND SOFTWARE AVAILABILITY

Processed data used for the analysis is available as supplementary data file Data S1. Raw “Illumina sequencing” data and the processed count data files that support the findings of this study have been deposited in NCBI's Gene Expression Omnibus and are accessible through GEO Series accession number GSE122806 (https://www.ncbi.nlm.nih.gov/geo/query/acc.cgi?acc=GSE122806) with reviewer token szgxsmygbxylhaz. Scripts are available from GitHub (https://github.com/lehner-lab/concentration_epistasis_CI).

## Supplemental tables

**Tables S1 – S8. Related to STAR Methods.**

## Supplemental Information titles

**Data S1.** Processed final data table for all single and double amino acid mutations analyzed in this study.

