## Supplementary figures for "Changes in gene expression shift and switch genetic interactions"

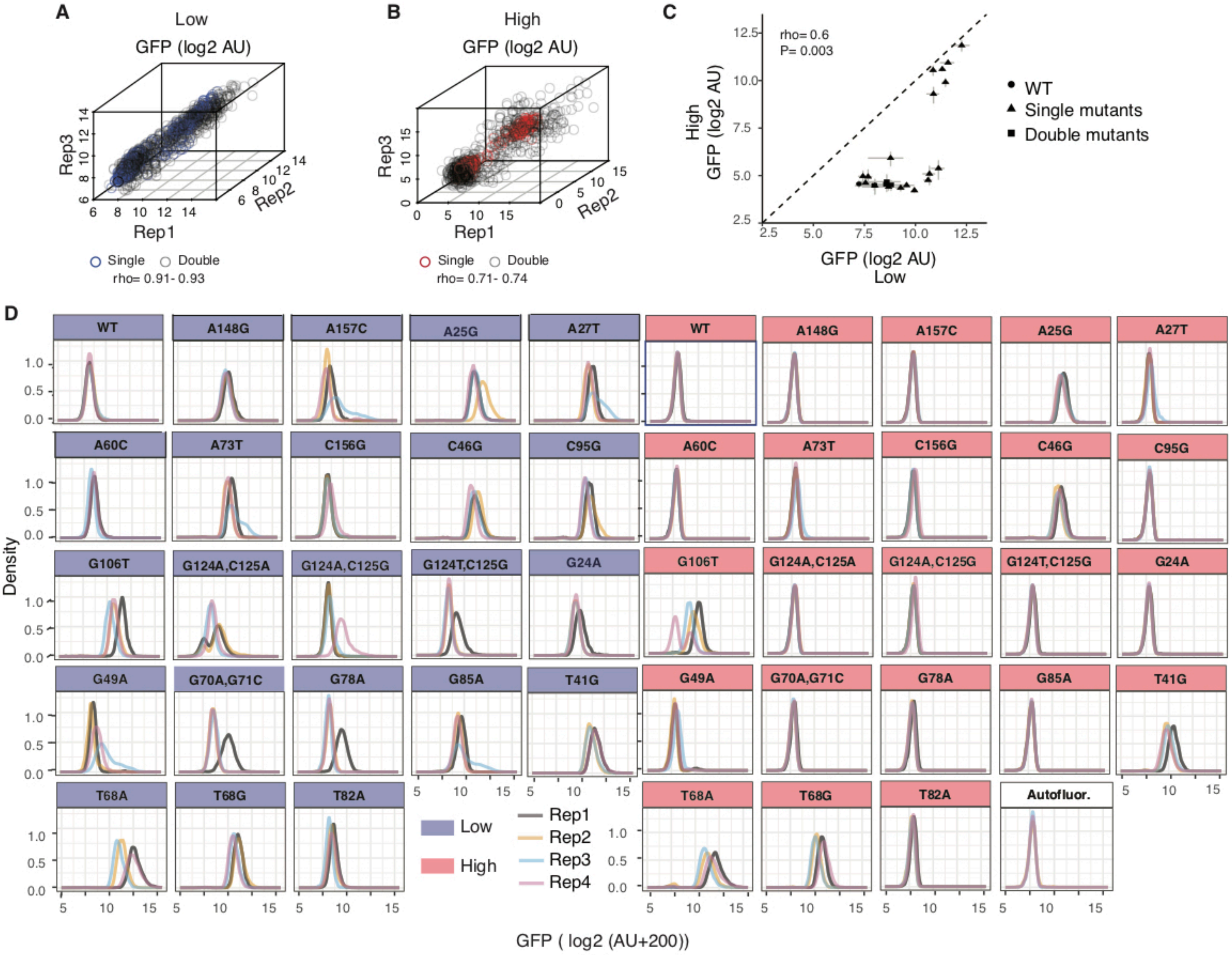

A
Low
GFP (log2 AU)
Rep3
Rep1
Rep2
Single
Double
rho= 0.91- 0.93
B
High
GFP (log2 AU)
Rep3
Rep1
Rep2
Single
Double
rho= 0.71- 0.74
C
rho= 0.6
P= 0.003
High
GFP (log2 AU)
GFP (log2 AU)
Low
WT
Single mutants
Double mutants
D
Density
WT
A148G
A157C
A25G
A27T
A60C
A73T
C156G
C46G
C95G
G106T
G124A,C125A
G124A,C125G
G124T,C125G
G24A
G49A
G70A,G71C
G78A
G85A
T41G
T68A
T68G
T82A
Autofluor.
Low
High
Rep1
Rep2
Rep3
Rep4
GFP ( log2 (AU+200))


Figure S1

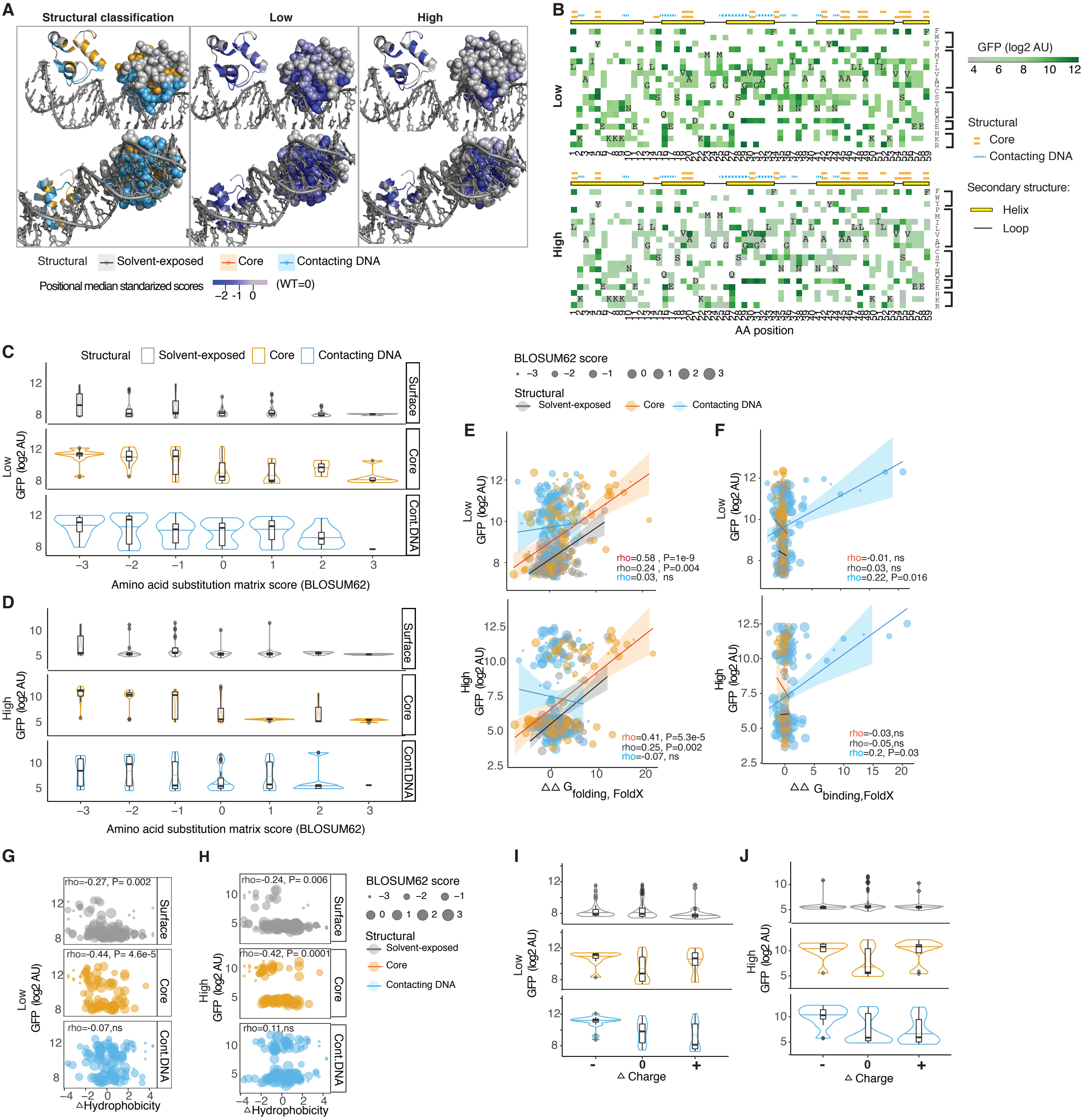
A
Structural classification
Low
High
Structural
Solvent-exposed
Core
Contacting DNA
Positional median standarized scores
-2 -1 0
(WT=0)
B
Low
High
GFP (log2 AU)
4 6 8 10 12
Structural
Core
Contacting DNA
Secondary structure:
Helix
Loop
AA position
C
Structural
Solvent-exposed
Core
Contacting DNA
Surface
Core
Cont.DNA
Low GFP (log2 AU)
Amino acid substitution matrix score (BLOSUM62)
D
High GFP (log2 AU)
Amino acid substitution matrix score (BLOSUM62)
BLOSUM62 score
-3 -2 -1 0 1 2 3
E
rho=0.58 , P=1e-9
rho=0.24 , P=0.004
rho=0.03, ns
rho=0.41, P=5.3e-5
rho=0.25, P=0.002
rho=-0.07, ns
ΔΔ G folding, FoldX
F
rho=-0.01, ns
rho=0.03, ns
rho=0.22, P=0.016
rho=-0.03,ns
rho=-0.05,ns
rho=0.2, P=0.03
ΔΔ G binding,FoldX
G
rho=-0.27, P= 0.002
rho=-0.44, P= 4.6e-5
rho=-0.07,ns
ΔHydrophobicity
H
rho=-0.24, P= 0.006
rho=-0.42, P= 0.0001
rho=0.11,ns
ΔHydrophobicity
I
Low GFP (log2 AU)
Δ Charge
J
High GFP (log2 AU)
Δ Charge

Figure S2

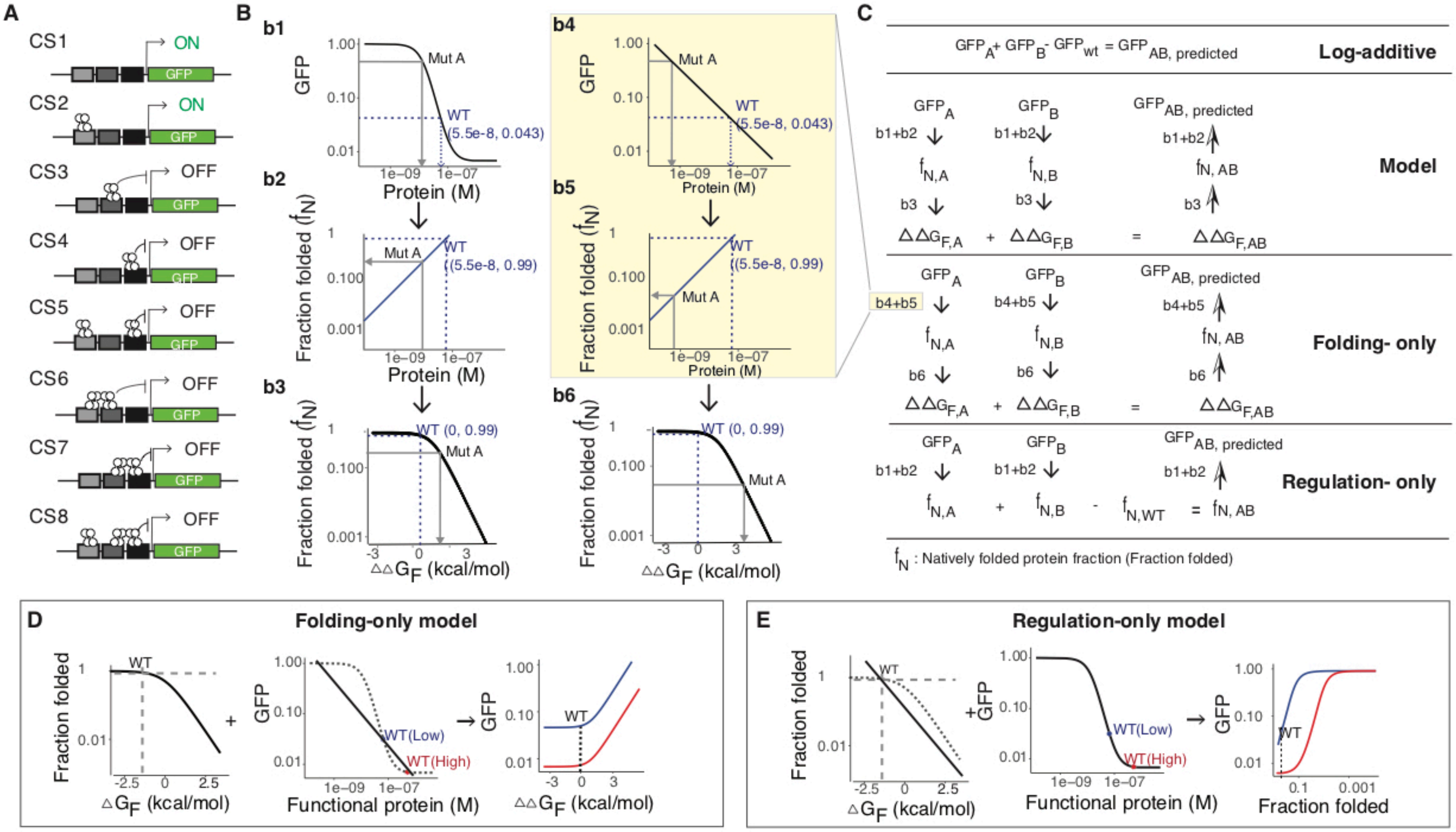
A
CS1
ON
CS2
ON
CS3
OFF
CS4
OFF
CS5
OFF
CS6
OFF
CS7
OFF
CS8
OFF
GFP
B
b1
b2
b3
b4
b5
b6
GFP
Fraction folded (fN)
Protein (M)
ΔΔGF (kcal/mol)
Mut A
WT
(5.5e-8, 0.043)
(5.5e-8, 0.99)
WT (0, 0.99)
C
GFPA+ GFPB- GFPwt = GFPAB, predicted
Log-additive
Model
Folding- only
Regulation- only
ΔΔGF,A + ΔΔGF,B = ΔΔGF,AB
fN,A + fN,B - fN,WT = fN, AB
fN : Natively folded protein fraction (Fraction folded)
D
Folding-only model
Fraction folded
ΔGF (kcal/mol)
Functional protein (M)
WT(Low)
WT(High)
ΔΔGF (kcal/mol)
E
Regulation-only model
Fraction folded

Figure S3

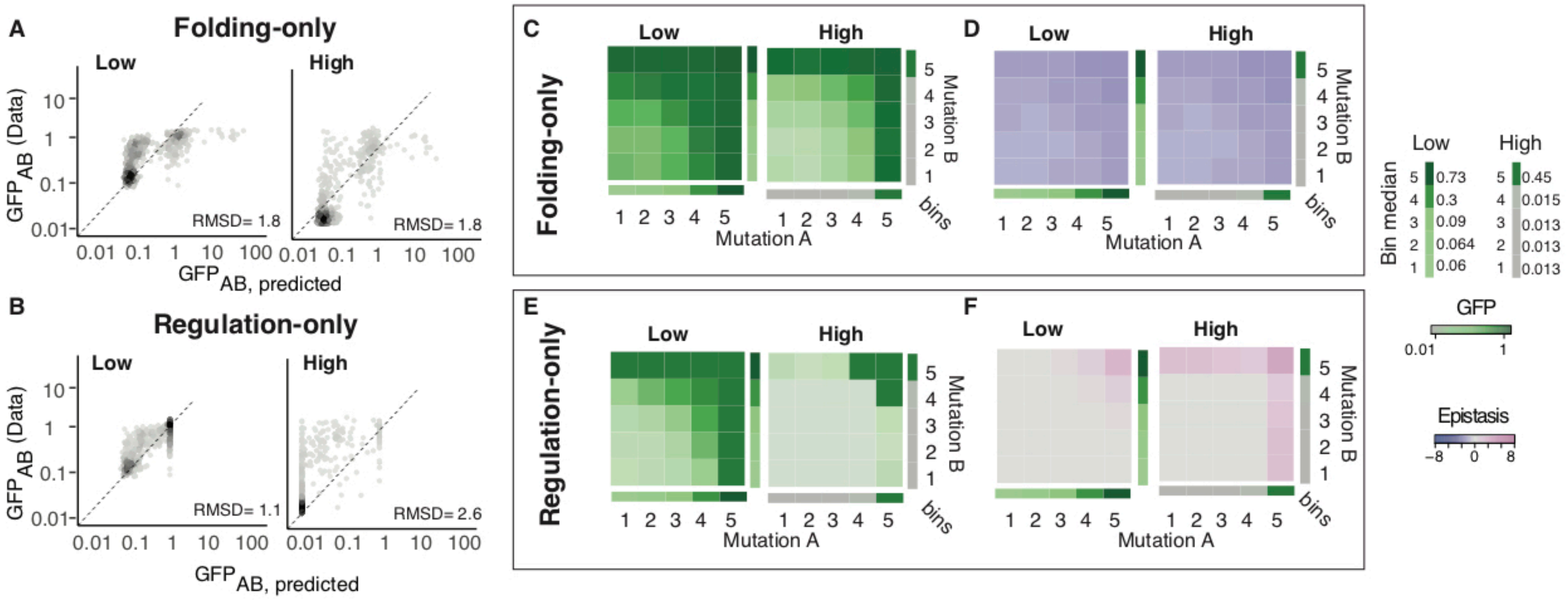

A
Folding-only
Low
High
GFPAB (Data)
GFPAB, predicted
RMSD= 1.8
RMSD= 1.8
B
Regulation-only
RMSD= 1.1
RMSD= 2.6
C
D
E
F
Mutation A
Mutation B
bins
Bin median
GFP
Epistasis


Figure S4

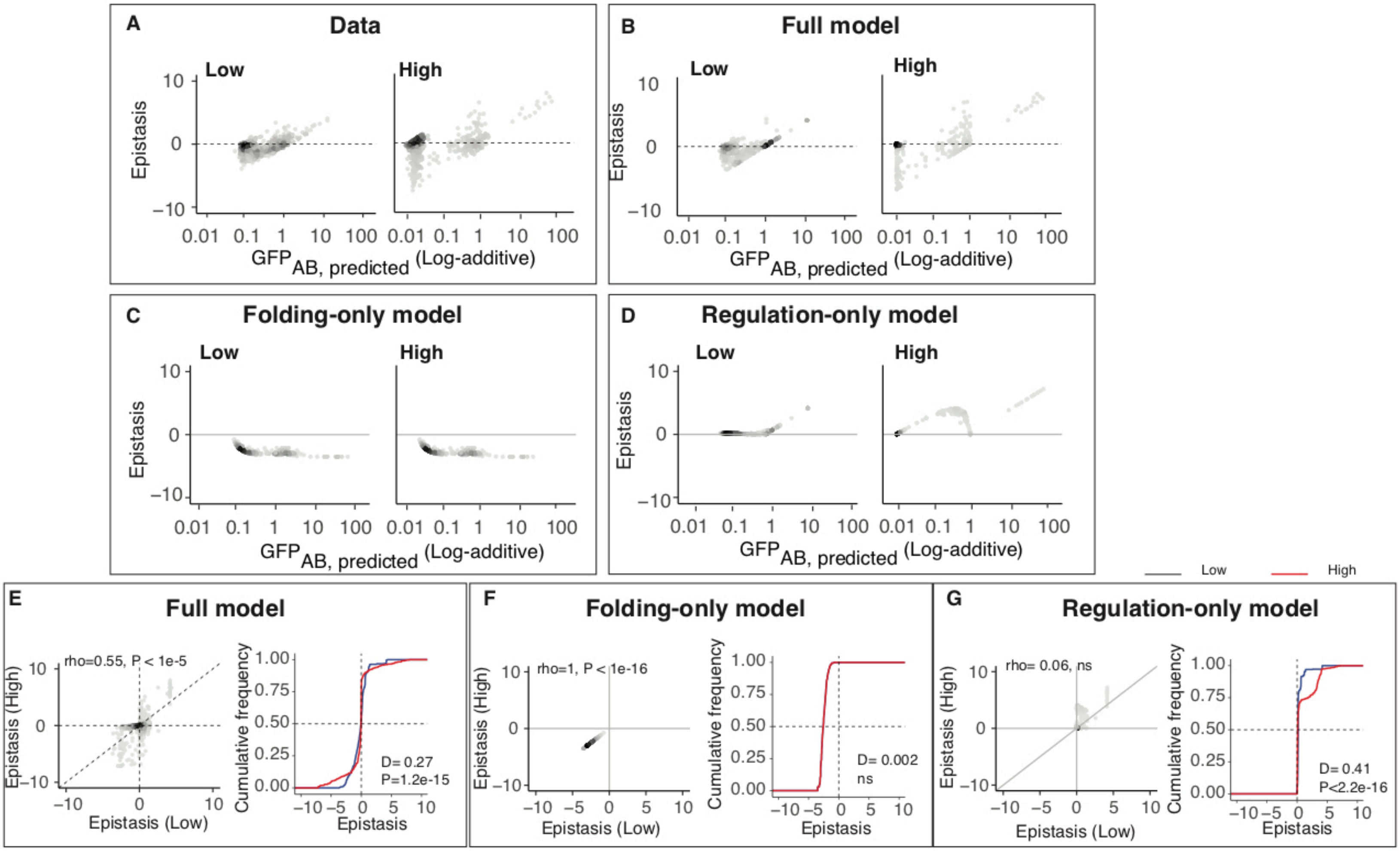

A
Data
Low
High
Epistasis
10
0
−10
0.01 0.1 1 10 100
GFP_AB, predicted (Log-additive)
B
Full model
Low
High
Epistasis
C
Folding-only model
Low
High
Epistasis
D
Regulation-only model
Low
High
Epistasis
Low
High
E
Full model
rho=0.55, P < 1e-5
Epistasis (High)
Epistasis (Low)
Cumulative frequency
1.00
0.75
0.50
0.25
0.00
D= 0.27
P=1.2e-15
Epistasis
F
Folding-only model
rho=1, P < 1e-16
Epistasis (High)
Epistasis (Low)
Cumulative frequency
D= 0.002
ns
Epistasis
G
Regulation-only model
rho= 0.06, ns
Epistasis (High)
Epistasis (Low)
Cumulative frequency
D= 0.41
P<2.2e-16
Epistasis


Figure S5

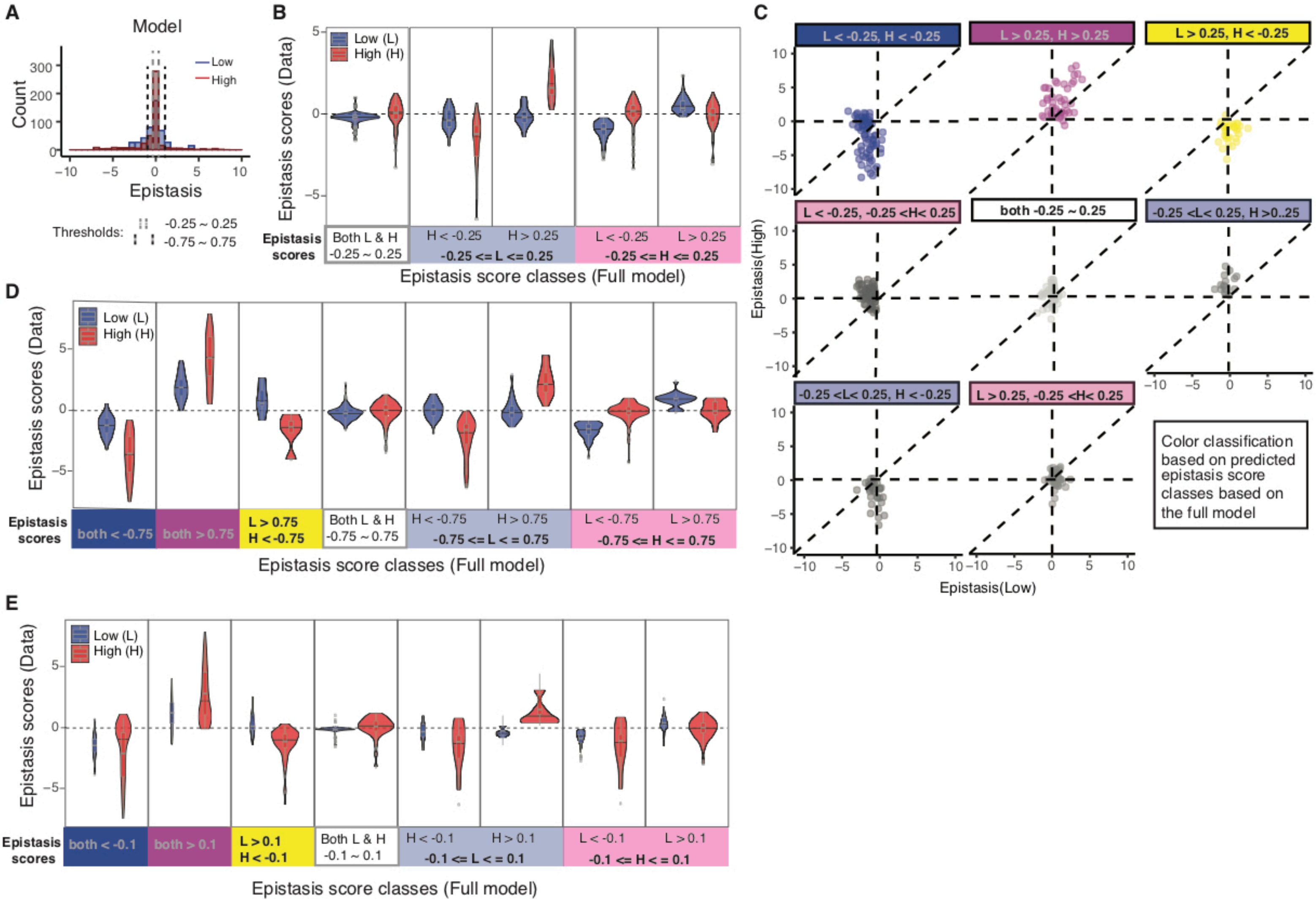
A
Model
Count
Epistasis
Low
High
Thresholds:
-0.25 ~ 0.25
-0.75 ~ 0.75
B
Epistasis scores (Data)
Low (L)
High (H)
Epistasis scores
Both L & H -0.25 ~ 0.25
H < -0.25
H > 0.25
-0.25 <= L <= 0.25
L < -0.25
L > 0.25
-0.25 <= H <= 0.25
Epistasis score classes (Full model)
C
L < -0.25, H < -0.25
L > 0.25, H > 0.25
L > 0.25, H < -0.25
L < -0.25, -0.25 <H< 0.25
both -0.25 ~ 0.25
-0.25 <L< 0.25, H >0..25
-0.25 <L< 0.25, H < -0.25
L > 0.25, -0.25 <H< 0.25
Color classification based on predicted epistasis score classes based on the full model
Epistasis(High)
Epistasis(Low)
D
Epistasis scores (Data)
Low (L)
High (H)
Epistasis scores
both < -0.75
both > 0.75
L > 0.75
H < -0.75
Both L & H -0.75 ~ 0.75
H < -0.75
H > 0.75
-0.75 <= L < = 0.75
L < -0.75
L > 0.75
-0.75 <= H < = 0.75
Epistasis score classes (Full model)
E
Epistasis scores (Data)
Low (L)
High (H)
Epistasis scores
both < -0.1
both > 0.1
L > 0.1
H < -0.1
Both L & H -0.1 ~ 0.1
H < -0.1
H > 0.1
-0.1 <= L < = 0.1
L < -0.1
L > 0.1
-0.1 <= H < = 0.1
Epistasis score classes (Full model)

Figure S6

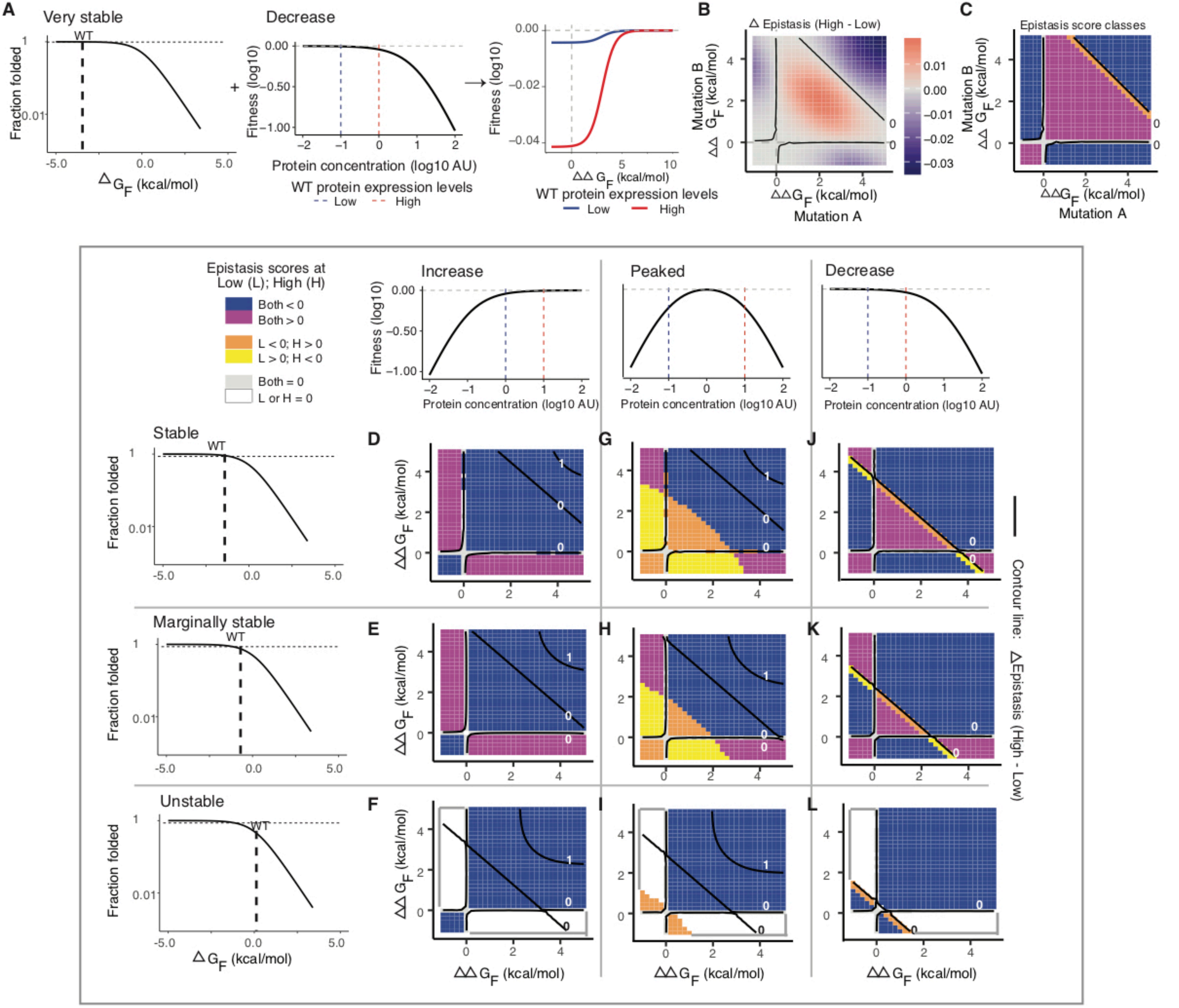
A
Very stable
Fraction folded
WT
ΔG_F (kcal/mol)
Decrease
Fitness (log10)
Protein concentration (log10 AU)
WT protein expression levels
Low
High
ΔΔG_F (kcal/mol)
B
Δ Epistasis (High - Low)
Mutation B ΔΔG_F (kcal/mol)
ΔΔG_F (kcal/mol)
Mutation A
C
Epistasis score classes
Epistasis scores at Low (L); High (H)
Both < 0
Both > 0
L < 0; H > 0
L > 0; H < 0
Both = 0
L or H = 0
Increase
Peaked
Decrease
Stable
Marginally stable
Unstable
D
E
F
G
H
I
J
K
L
Contour line: ΔEpistasis (High - Low)

Figure S7

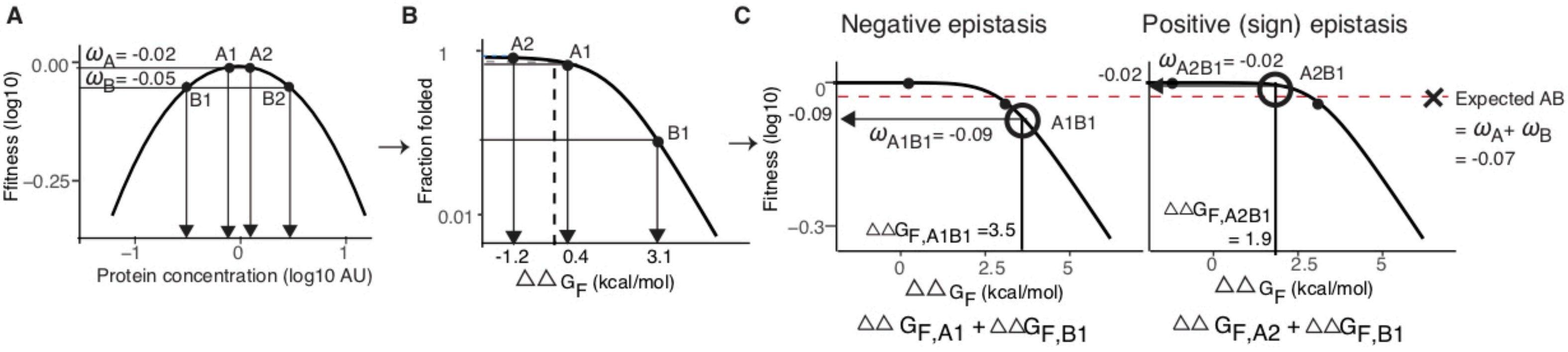
A
$\omega_A$= -0.02
$\omega_B$= -0.05
A1
A2
B1
B2
0.00
-0.25
Ffitness (log10)
-1
0
1
Protein concentration (log10 AU)
B
A2
A1
B1
1
0.01
Fraction folded
-1.2
0.4
3.1
$\Delta\Delta G_F$ (kcal/mol)
C
Negative epistasis
Positive (sign) epistasis
0
-0.09
-0.3
Fitness (log10)
$\omega_{A1B1}$= -0.09
A1B1
$\Delta\Delta G_{F,A1B1}$ =3.5
0
2.5
5
$\Delta\Delta G_F$ (kcal/mol)
$\Delta\Delta G_{F,A1} + \Delta\Delta G_{F,B1}$
-0.02
$\omega_{A2B1}$= -0.02
A2B1
$\Delta\Delta G_{F,A2B1}$
= 1.9
Expected AB
= $\omega_A$+ $\omega_B$
= -0.07
$\Delta\Delta G_{F,A2} + \Delta\Delta G_{F,B1}$

Figure S8

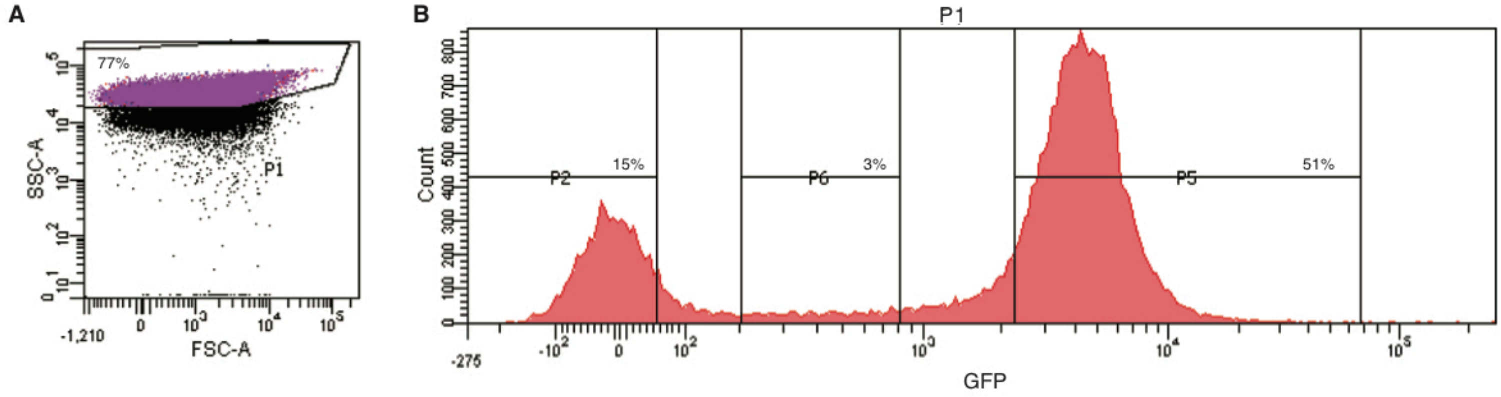


C

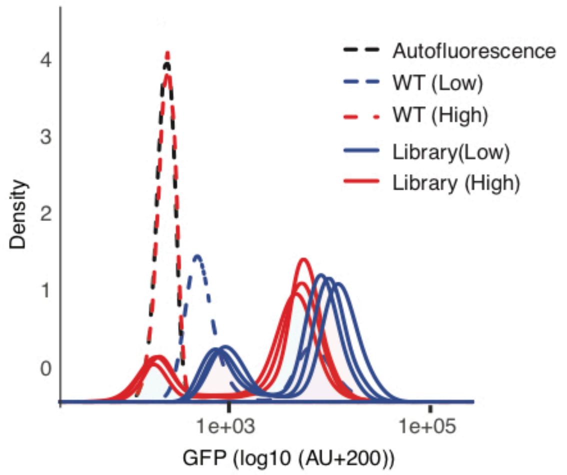


$$\text{GFP\_index} = \frac{\text{Mean (No\_repression)- Mean (wt\_repression)}}{\text{2 X SD (wt\_repression)}}$$

**GFP_index**

| | replicate 1 | replicate 2 | replicate 3 |
|---|---|---|---|
| Library (Low) | 0.69 | 0.68 | 0.75 |
| Library (High) | 1763 | 1974 | 924 |

Figure S9

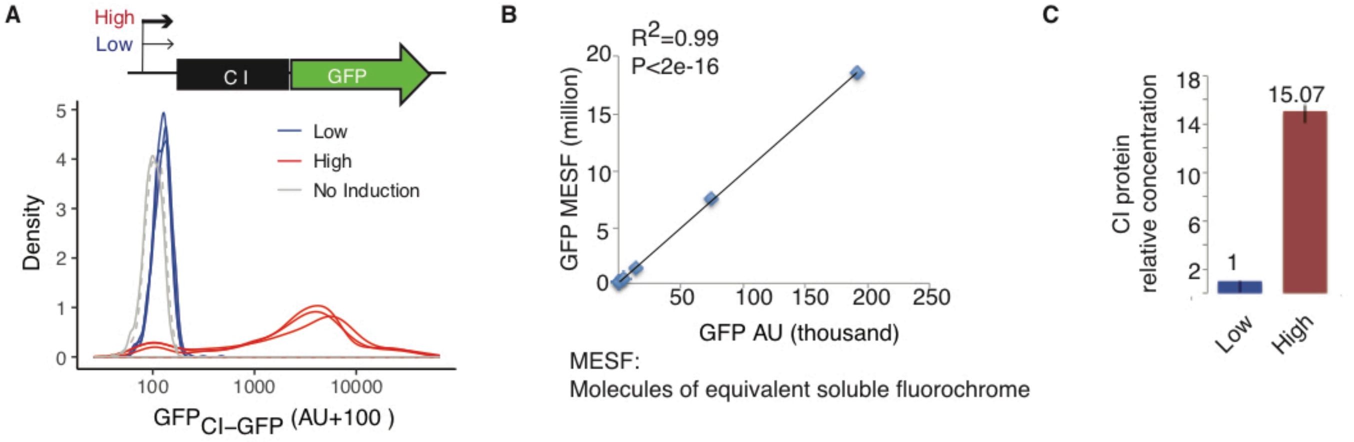
A
High
Low
C I
GFP
Density
Low
High
No Induction
GFP CI–GFP (AU+100 )
B
R2=0.99
P<2e-16
GFP MESF (million)
GFP AU (thousand)
MESF:
Molecules of equivalent soluble fluorochrome
C
CI protein
relative concentration
1
15.07
Low
High

Figure S10

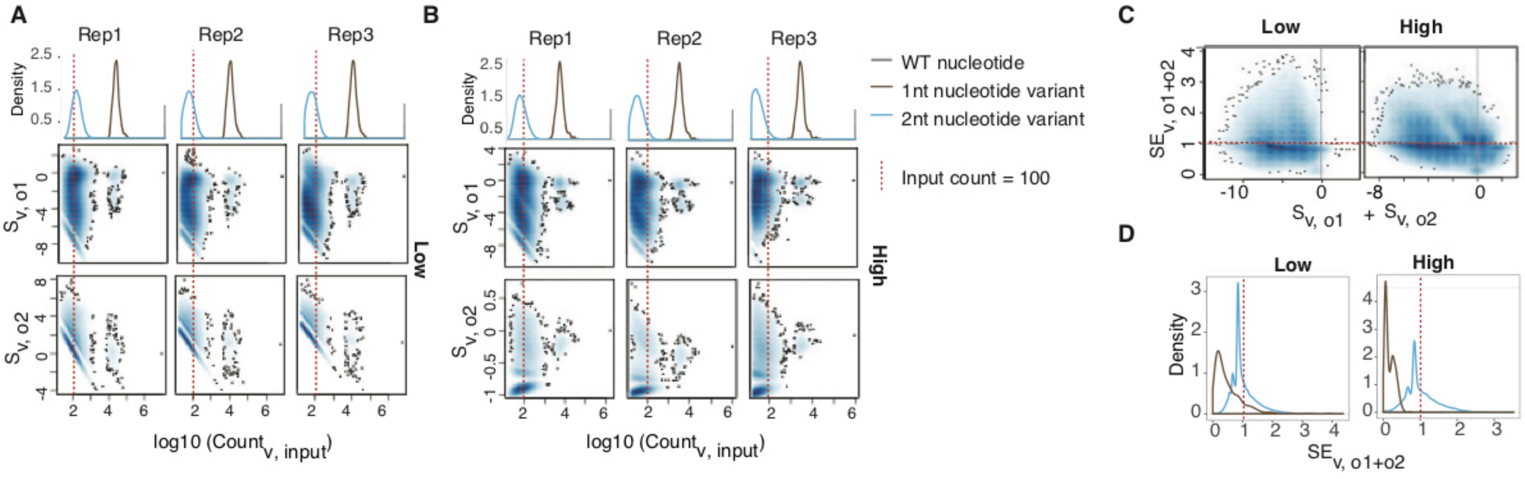
A
Rep1
Rep2
Rep3
Density
Low
log10 (Count_v, input)
B
High
WT nucleotide
1nt nucleotide variant
2nt nucleotide variant
Input count = 100
C
Low
High
D
Low
High
Density

Figure S11

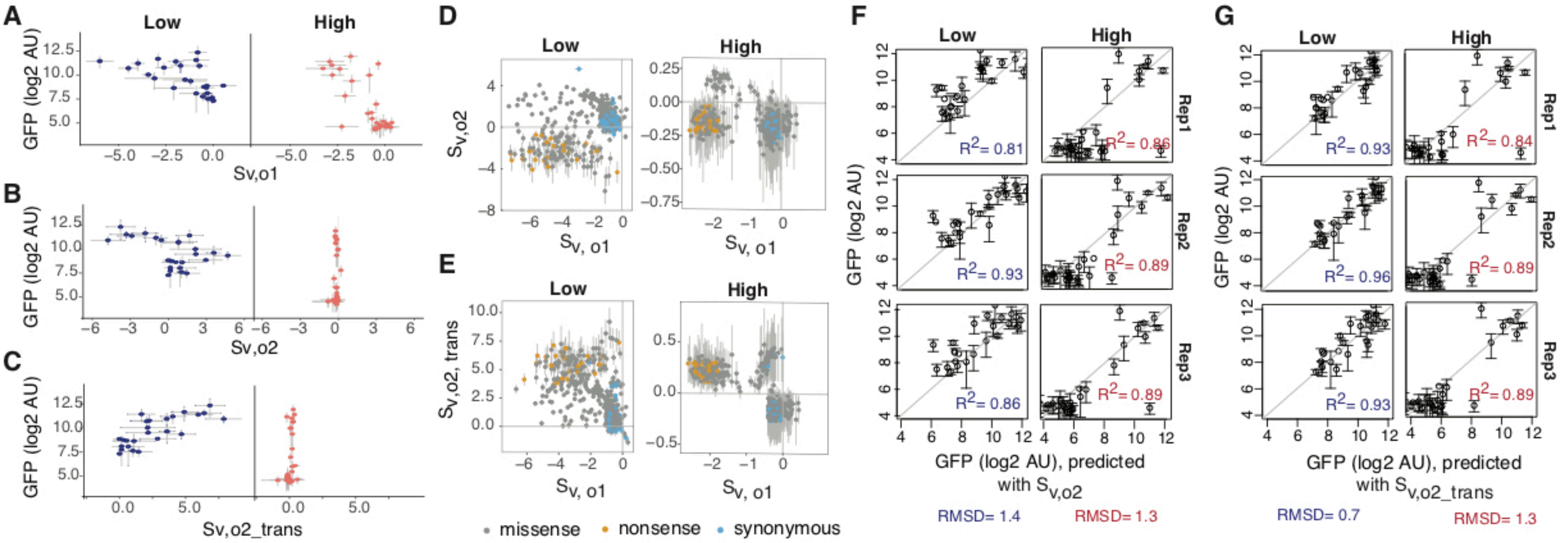
A
B
C
D
E
F
G
Low
High
GFP (log2 AU)
Sv,o1
Sv,o2
Sv,o2_trans
$S_{v,o2}$
$S_{v, o1}$
$S_{v,o2, trans}$
missense
nonsense
synonymous
Rep1
Rep2
Rep3
$R^2$= 0.81
$R^2$= 0.86
$R^2$= 0.93
$R^2$= 0.89
$R^2$= 0.86
$R^2$= 0.89
$R^2$= 0.93
$R^2$= 0.84
$R^2$= 0.96
$R^2$= 0.89
$R^2$= 0.93
$R^2$= 0.89
GFP (log2 AU), predicted with $S_{v,o2}$
GFP (log2 AU), predicted with $S_{v,o2\_trans}$
RMSD= 1.4
RMSD= 1.3
RMSD= 0.7
RMSD= 1.3

Figure S12

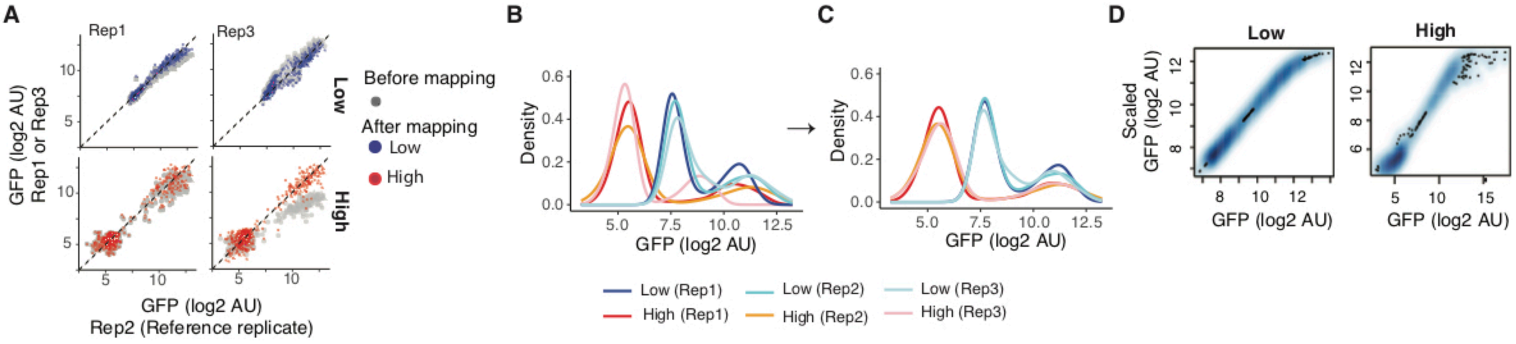
A
Rep1
Rep3
Low
High
Before mapping
After mapping
Low
High
GFP (log2 AU)
Rep1 or Rep3
GFP (log2 AU)
Rep2 (Reference replicate)
B
C
Density
GFP (log2 AU)
Low (Rep1)
High (Rep1)
Low (Rep2)
High (Rep2)
Low (Rep3)
High (Rep3)
D
Low
High
Scaled
GFP (log2 AU)
GFP (log2 AU)

Figure S13

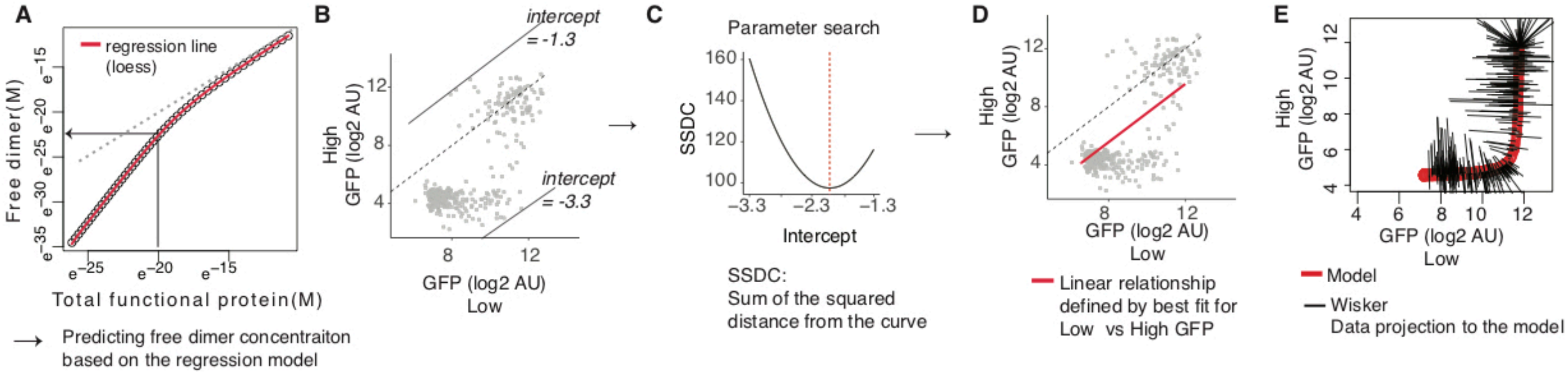

A
regression line
(loess)
Free dimer(M)
Total functional protein(M)
Predicting free dimer concentraiton
based on the regression model
B
intercept
= -1.3
intercept
= -3.3
High
GFP (log2 AU)
GFP (log2 AU)
Low
C
Parameter search
SSDC
Intercept
SSDC:
Sum of the squared
distance from the curve
D
High
GFP (log2 AU)
GFP (log2 AU)
Low
Linear relationship
defined by best fit for
Low vs High GFP
E
High
GFP (log2 AU)
GFP (log2 AU)
Low
Model
Wisker
Data projection to the model


Figure S14
