## Supplementary tables for "Changes in gene expression shift and switch genetic interactions"

**Table S1**. PCR primers

| F: Forward; R: Reverse, 5' to 3' | | Note |
|---|---|---|
| d_CIF | ACACAAGAGCAGCTTGAGGA | "doped" library amplification |
| d_CIR | ATTTCTCTGGCGATTGAAGG | |
| pBAD-CI-F | TCCTCAAGCTGCTCTTGTGT | For Gibson Assemble CI to the backbone |
| pBAD-CI-R | CCTTCAATCGCCAGAGAAAT | |
| Q5SDM_CIF | ATGAGCACAAAAAAGAAACC | Changing 5prime UTR for expression of CI |
| Q5SDM_CIR | GGTTAATTCCTCCTGTTAG | |
| CI_GFP_fuse_F | GCTGGTTCTGGCGAATTCATGCGTAAAGGCGAAGAAC | Making CI-GFP fusion for protein quantification |
| CI_GFP_fuse_R | AGCGGAGCCAGCGGATCCGCCAAACGTCTCTTCAGG | |
| colony-F1 | GGCGTCACACTTTGCTATGC | For colony PCR |
| colony-R1 | ACAGTTCTTCGCCTTTACGC | |
| d_CI_F_s | GCTTGAGGACGCACGTC | "doped" library amplification for sequencing library |
| d_CI_R_s | TCTGGCGATTGAAGGGCT | |
| d_CI_Fs_1 | GAACGTTCGCTTGAGGACGCACGTC | Input.rep1.barcoded |
| d_CI_Rs_1 | GAACGTTCTCTGGCGATTGAAGGGCT | |
| d_CI_Fs_2 | GCCGAATTGCTTGAGGACGCACGTC | Input.rep2.barcoded |
| d_CI_Rs_2 | GCCGAATTTCTGGCGATTGAAGGGCT | |
| d_CI_Fs_3 | CGGCAATTGCTTGAGGACGCACGTC | Input.rep3.barcoded |
| d_CI_Rs_3 | CGGCAATTTCTGGCGATTGAAGGGCT | |
| d_CI_Fs_4 | GCGCATATGCTTGAGGACGCACGTC | Output1 &2.rep1.barcoded |
| d_CI_Rs_4 | GCGCATATTCTGGCGATTGAAGGGCT | |
| d_CI_Fs_5 | CAACCATGGCTTGAGGACGCACGTC | Output1 &2.rep2.barcoded |
| d_CI_Rs_5 | CAACCATGTCTGGCGATTGAAGGGCT | |
| d_CI_Fs_6 | CGTACCTTGCTTGAGGACGCACGTC | Output1 &2.rep3.barcoded |
| d_CI_Rs_6 | CGTACCTTTCTGGCGATTGAAGGGCT | |
| Q5SDM1_F | TTGATGCCATTAAATAAAGCAC | G124T,C125G |
| Q5SDM1.1_R | TGCATTAAATcaTTATAACGCCGCATTG | |
| Q5SDM1_F | TTGATGCCATTAAATAAAGCAC | G124A,C125G |
| Q5SDM1.2_R | TGCATTAAATctTTATAACGCCGCATTG | |
| Q5SDM1_F | TTGATGCCATTAAATAAAGCAC | G124A,C125A |
| Q5SDM1.3_R | TGCATTAAATttTTATAACGCCGCATTG | |
| Q5SDM4_F | AGACAAGATGacGATGGGGCAGTC | G70A,G71C |
| Q5SDM4_R | GCGACAGATTCCTGGGAT | |
| Q5SDM5_F | AGGAATCTGTCGCcGACAAGATGGGGA | A60C (synonymous) |
| Q5SDM5_R | TCCCCATCTTGTCGGCGACAGATTCCT | |
| Q5SDM6_F | GCAATTTATGAAAAAAAaAAAAATGAACTTGGCTT | G78A (synonymous) |
| Q5SDM6_R | AAGCCAAGTTCATTTTTTTTTTTTTCATAAATTGC | |
| Q5SDM7_F | CTTGCAAAAATTCTgAAAGTTAGCGTTGAAGAATTTAGC | C156G(synonymous) |
| Q5SDM7_R | GCTAAATTCTTCAACGCTAACTTTCAGAATTTTTGCAAG | |
| Q5SDM8_F | AAATGAACTTGGCTgATCCCAGGAATCTGTCG | T41G (nonsense) |
| Q5SDM8_R | CGACAGATTCCTGGGATCAGCCAAGTTCATTT | |
| Q5SDM9_F | CTTATCCCAGaAATCTGTCGC | G49A |
| Q5SDM9_R | CCAAGTTCATTTTTCTTTTTTTC | |
| Q5SDM10_F | GCAATTTATGAAAAAAAaAAAAATGAACTTGGCTT | G24A |
| Q5SDM10_R | AAGCCAAGTTCATTTTTTTTTTTTTCATAAATTGC | |
| Q5SDM11_F | TCGCAGACAAGAaGGGGATGGGGCAG | T68A |
| Q5SDM11_R | CTGCCCCATCCCCtTCTTGTCTGCGA | |
| Q5SDM12_F | AAAAAAAGAAtAATGAACTTGGCTTATC | A27T |
| Q5SDM12_R | CATAAATTGCTTTAAGGCG | |
| Q5SDM13_F | TGAAAAAAAGGAAAATGAACTTGG | A25G |
| Q5SDM13_R | TAAATTGCTTTAAGGCGAC | |
| Q5SDM14_F | AAAAATTCTCCAAGTTAGCGTTGAAGAATTTAGC | A157C |
| Q5SDM14_R | GCAAGCAATGCGGCGTTA | |
| Q5SDM15_F | GGCGTTGGTGgTTTATTTAATGGC | C95G |
| Q5SDM15_R | TGACTGCCCCATCCCCAT | |
| Q5SDM16_F | GGGGCAGTCAaGCGTTGGTGC | G85A |
| Q5SDM16_R | ATCCCCATCTTGTCTGCGACAG | |
| Q5SDM17_F | ATTGCTTGCAgAAATTCTCAAAGTTAG | A148G |
| Q5SDM17_R | GCGGCGTTATAAGCATTTAATG | |
| Q5SDM18_F | TTTATTTAATtGCATCAATGCATTAAATGCTTATAACGCC | G106T |
| Q5SDM18_R | GCACCAACGCCTGACTGC | |
| Q5SDM19_F | CAAGATGGGGtTGGGGCAGTC | A73T |
| Q5SDM19_R | TCTGCGACAGATTCCTGG | |
| Q5SDM20_F | TGGCTTATCCGAGGAATCTGTC | C46G |
| Q5SDM20_R | AGTTCATTTTTCTTTTTTTCATAAATTG | |
| Q5SDM21_F | GCAGACAAGAgGGGGATGGGG | T68G |
| Q5SDM21_R | GACAGATTCCTGGGATAAGCC | |
| Q5SDM22_F | GATGGGGCAGaCAGGCGTTGG | T82A |
| Q5SDM22_R | CCCATCTTGTCTGCGACAG | |

Lower case letter in the primer sequences indicate the targeted mutation to be incorporated.

**Table S2**. Individually validated confirmation set

| ID | category | WT | Substitution | Position |
|---|---|---|---|---|
| G24A | missense | G | A | 24 |
| A25G | missense | A | G | 25 |
| A27T | missense | A | T | 27 |
| T41G | nonsense | T | G | 41 |
| C46G | missense | C | G | 46 |
| G49A | missense | G | A | 49 |
| A60C | synonymous | A | C | 60 |
| T68G | missense | T | G | 68 |
| A73T | missense | A | T | 73 |
| G78A | synonymous | G | A | 78 |
| T68A | missense | T | A | 78 |
| T82A | missense | T | A | 82 |
| G85A | missense | G | A | 85 |
| C95G | missense | C | G | 95 |
| G106T | missense | G | T | 106 |
| A148G | missense | A | G | 148 |
| C156G | synonymous | C | G | 156 |
| A157C | missense | A | C | 157 |
| G70A,G71C | missense | G,G | A,C | 70,71 |
| G124T,C125G | missense | G,C | T,G | 124,125 |
| G124A,C125G | missense | G,C | A,G | 124,125 |
| G124A,C125A | missense | G,C | A,A | 124,125 |

**Table S3.** Coefficients for linear models to predict GFP signals from enrichment scores

| Expression | Intercept<br>α | Sv,o1<br>β | Sv,o2_trans<br>γ | adj- $R^2$ | P |
|---|---|---|---|---|---|
| Low | 7.23*** | -0.51*** | 0.52*** | 0.96 | 5.1e-15 |
| High | 4.56*** | -2.23*** | -1.64 | 0.84 | 3.6e-9 |

Significance code P<1e-4 ***

**Table S4**. Covariance of Enrichment scores Sv,o1 and Sv,o2,trans

| Expression level | Rep1 | Rep2 | Rep3 |
|---|---|---|---|
| Low | -1.48 | -1.95 | -1.85 |
| High | -0.16 | -0.22 | -0.12 |

**Table S5**. Coefficients to map replicates 1 and 3 to replicate2

| | α1 | β1 | α3 | β3 |
|---|---|---|---|---|
| Low | 1.5±0.02 | 0.9±0.02 | 1.3±0.02 | 0.9±0.002 |
| High | 1.7±0.03 | 0.8±0.03 | 0.7±0.04 | 1.1±0.05 |

**Table S6**. Configuration states and the energy terms from Ackers' model

| CS*i* | Occupied OR | CI dimer (N*i*) | Downstream gene | Total energy ($\Delta G_{CS}$) |
|---|---|---|---|---|
| 1 | − | 0 | ON | 0 |
| 2 | OR3 | 1 | ON | $\Delta G_3$ |
| 3 | OR2 | 1 | OFF | $\Delta G_2$ |
| 4 | OR1 | 1 | OFF | $\Delta G_1$ |
| 5 | OR2, OR3 | 2 | OFF | $\Delta G_2 + \Delta G_3 + \Delta G_{co}$ |
| 6 | OR1, OR2 | 2 | OFF | $\Delta G_1 + \Delta G_2 + \Delta G_{co}$ |
| 7 | OR1, OR3 | 2 | OFF | $\Delta G_1 + \Delta G_3$ |
| 8 | OR1, OR2, OR3 | 3 | OFF | $\Delta G_1 + \Delta G_2 + \Delta G_3 + \Delta G_{co}$ |

**Table S7**. Parameters for CI regulatory interaction model, from the literature

| | |
|---|---|
| $K_a$ | $5\times10^7$ |
| [OR] | $10^{-9}$ M |
| $\Delta G_1$ | -11.7 kcal |
| $\Delta G_2$ | -10.1 kcal |
| $\Delta G_3$ | -10.1 kcal |
| $\Delta G_{co}$ | -2 kcal |

**Table S8**. Parameters estimated for modeling transcription regulatory interaction

| | | |
|---|---|---|
| $[CI_{E,low}]$ | $5.5\times10^{-8}$ M | Calculated based on $GFP_{wt,low}$ |
| $[CI_{E,high}]$ | $8.4\times10\text{-}^{7}$ M | Calculated based on $GFP_{wt,high}$ |
